# Probe-Seq enables transcriptional profiling of specific cell types from heterogeneous tissue by RNA-based isolation

**DOI:** 10.1101/735738

**Authors:** Ryoji Amamoto, Mauricio D. Garcia, Emma R. West, Jiho Choi, Sylvain W. Lapan, Elizabeth A. Lane, Norbert Perrimon, Constance L. Cepko

## Abstract

Recent transcriptional profiling technologies are uncovering previously-undefined cell populations and molecular markers at an unprecedented pace. While single cell RNA (scRNA) sequencing is an attractive approach for unbiased transcriptional profiling of all cell types, a complementary method to isolate and sequence specific cell populations from heterogeneous tissue remains challenging. Here, we developed Probe-Seq, which allows deep transcriptional profiling of specific cell types isolated using RNA as the defining feature. Dissociated cells are labelled using fluorescent *in situ* hybridization (FISH) for RNA, and then isolated by fluorescent activated cell sorting (FACS). We used Probe-Seq to purify and profile specific cell types from mouse, human, and chick retinas, as well as the *Drosophila* midgut. Probe-Seq is compatible with frozen nuclei, making cell types within archival tissue immediately accessible. As it can be multiplexed, combinations of markers can be used to create specificity. Multiplexing also allows for the isolation of multiple cell types from one cell preparation. Probe-Seq should enable RNA profiling of specific cell types from any organism.

## INTRODUCTION

Multicellular eukaryotic tissues often comprise many different cell types, commonly classified using their morphological features, physiological function, anatomical location, and/or molecular markers. For example, the retina, a thin sheet of neural tissue in the eye that transmits visual information to the brain, contains seven major cell classes - rods, cones, bipolar cells (BC), amacrine cells (AC), horizontal cells (HC), Müller glia (MG), and retinal ganglion cells (RGC), first defined primarily using morphology^1,2^. More recently, scRNA profiling technologies have led to the appreciation of many subtypes of these major cell classes, bringing the total number of retinal cell types close to 100^3–5^. Such accelerated discovery of cellular diversity is not unique to the retina, as scRNA profiling is being carried out in many tissues and organisms^6–8^.

Several approaches have been used to transcriptionally profile tissues. Bulk RNA sequencing of whole tissues can be done at great depth, but does not capture the diversity of individual transcriptomes and often fails to reflect signatures of rare cell types. Currently, bulk sequencing of specific cell types is limited by the availability of cell type-specific promoters, enhancers, dyes, or antigens for their isolation^9–14^. This has limited bulk RNA sequencing primarily to select cell types in genetically-tractable organisms. Single cell and single nucleus RNA sequencing methods have allowed for the recording of transcriptional states of many individual cells simultaneously^3,5,15–18^. Despite the undeniable appeal of scRNA sequencing, capturing deep profiles of specific cell populations in bulk can be sufficient or preferable for many experiments, e.g. when the goal is to understand the results of perturbations.

We and others have used antibodies to enable FACS-based isolation for transcriptional profiling of specific cell populations^11,19–23^. However, antibodies are frequently unavailable for a specific cell type. Furthermore, marker proteins in certain cell types such as neurons are often localized to processes that are lost during cellular dissociation. We therefore aimed to create a method that would leverage the newly discovered RNA expression patterns for the isolation of specific cell populations from any organism. This led us to develop Probe-Seq, which uses a FISH method based upon a new probe design, Serial Amplification By Exchange Reaction (SABER)^24^. Probe-Seq uses RNA markers expressed in specific cell types to label cells for isolation by FACS and subsequent transcriptional profiling. Although specific cells cultured *in vitro* have been successfully labelled by FISH for isolation using FACS, this method had not yet been tested for tissue^25–27^. We used Probe-Seq to isolate rare bipolar cells from the mouse retina, cell types that were previously defined using scRNA sequencing^5^. We demonstrate that probe sets for multiple genes can be hybridized at once, allowing isolation of multiple cell types simultaneously. Moreover, the fluorescent oligonucleotides used to detect the probe sets can be quickly hybridized and then stripped. This enables isolation of an indefinite number of cell types from one sample by serial sorting and re-labeling. We extended Probe-Seq to specific bipolar cell subtypes in frozen archival human retina by labeling nuclear RNA. To further test the utility of Probe-Seq in non-vertebrate animals and non-CNS tissues, we profiled intestinal stem cells from the *Drosophila* gut. In each of these experiments, the transcriptional profiles of isolated populations closely matched those obtained by scRNA sequencing, and in most cases, the number of genes detected exceeded 10,000. Finally, we used Probe-Seq on the chick retina, an organism that is difficult to genetically manipulate, to determine the transcriptional profile of a subset of developing retinal cells that give rise to the chick high acuity area. Taken together, Probe-Seq is a method that enables deep transcriptional profiling of specific cell types in heterogeneous tissue from potentially any organism.

## RESULTS

### Specific bipolar cell subtypes can be isolated and profiled from the mouse retina using Probe-Seq

To determine whether Probe-Seq can enable the isolation and profiling of specific cell types based on FISH labeling, we tested it using the mouse retina. The retina is a highly heterogeneous tissue, with cell classes and subtypes classified by scRNA sequencing, as well as more classical methods^2^. We used a new method for FISH, SABER-FISH, to label the intracellular RNA^24^. SABER-FISH uses OligoMiner to design 20-40 nt tiling oligonucleotides that are complementary to the RNA species of interest and are optimized for minimal off-target binding. The tiling oligonucleotides are pooled to generate a gene-specific probe set. Each probe set is then extended using a primer exchange reaction^28^, which appends many copies of a short-repeated sequence to each tiling oligonucleotide in the set. These concatemer sequences can be made unique for each probe set. The concatemers are detected by the hybridization of short fluorescent oligonucleotides.

To isolate specific BC subtypes, fresh adult mouse retinas were dissociated, fixed, and permeabilized prior to FISH labeling (Figure 1a). We designed gene-specific probe sets against *Vsx2*, a marker of all BCs and MG, and *Grik1*, a marker of BC2, BC3A, BC3B, and BC4 subtypes (~2% of all retinal cells) (Figure 1b). *Vsx2* and *Grik1* probe sets were hybridized to the dissociated retinal cells overnight at 43°C, and fluorescent oligonucleotides were subsequently hybridized to the gene-specific probe sets. By FACS, single cells were identified by gating for a single peak of Hoechst^+^ events, while debris and doublets were excluded (Figure 1c). Out of these single cells, the *Vsx2*^+^ population was judged to be the cells that shifted away from the diagonal *Vsx2*^−^ events (Figure 1c). Out of the *Vsx2*^+^ population, we found three populations that were *Grik1*^−^, *Grik1*^MID^, and *Grik1*^HI^. Based upon scRNA sequencing of BC subtypes, *Grik1*^MID^ likely corresponded to BC2, and *Grik1*^HI^ to BC3A, BC3B, and BC4^5^. We isolated both *Grik1*^MID^ and *Grik1*^HI^ (henceforth called *Grik1*^+^) cells as well as *Vsx2*^+^/*Grik1*^−^ (henceforth called *Grik1*^−^) and *Vsx2*^−^ cell populations. We expected the *Grik1*^+^ population to contain BC2-BC4, the *Grik1*^−^ population to contain other BC subtypes and MG, and the *Vsx2*^−^ population to contain non-BC/MG cell types (Figure 1d). The isolated populations displayed the expected FISH puncta (Figure 1e). On average, we isolated 200,000±0 *Vsx2*^−^ cells, 96,000±18,600 *Grik1*^−^ cells, and 22,000±1,000 *Grik1*^+^ cells per biological replicate, with each replicate originating from 2 retinas. These results indicate that gene-specific SABER-FISH can label dissociated cell populations for isolation by FACS.

**Figure 1:**
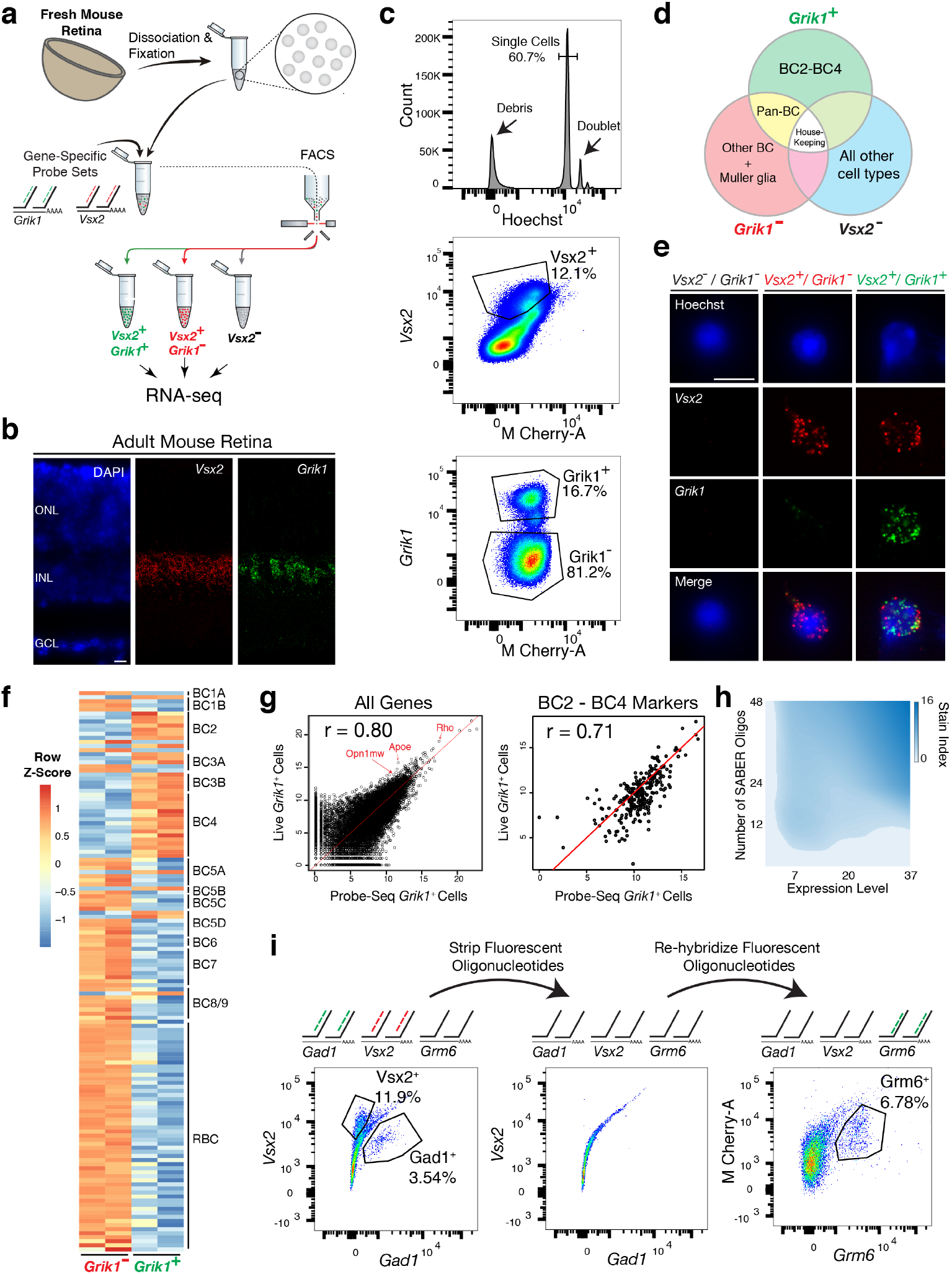
Isolation and transcriptional profiling of specific BC subtypes from the adult mouse retina. (**a**) Schematic of Probe-Seq for the adult mouse retina. The retina was dissociated into single cells, fixed, and permeabilized. Cells were incubated with gene-specific probe sets against *Vsx2* and *Grik1* and subsequently incubated with fluorescent oligonucleotides. Three populations of labeled cells (*Vsx2*^−^, *Vsx2*^+^/*Grik1*^−^, and *Vsx2*^+^/*Grik1*^+^) were isolated by FACS for downstream RNA sequencing. (**b**) SABER FISH signals from an adult mouse retina section using *Vsx2* and *Grik1* probe sets. (**c**) Representative FACS plot of all events (top panel) on a Hoechst histogram. The debris is the peak near 0. The first peak after the debris is the single cell 2N peak. 4N doublets and other cell clumps are in the peaks to the right of the single cell peak. Representative FACS plots of all single cells (middle panel) with *Vsx2* fluorescence on the y-axis and empty M-Cherry-A autofluorescence on the x-axis. The negative population ran along the diagonal. The *Vsx2*^+^ population (12.1%) was left shifted, indicating high *Vsx2* fluorescence and low M-Cherry-A autofluorescence. FACS plot of only the *Vsx2*^+^ population (bottom panel) with *Grik1* fluorescence on the y-axis and empty M-Cherry-A autofluorescence on the x-axis. *Vsx2*^+^/*Grik1*^+^ population (16.7%) displayed strong separation from the *Vsx2*^+^/*Grik1*^−^ population (81.2%). The *Grik1*^MID^ population was included in the *Vsx2*^+^/*Grik1*^+^ population. (**d**) Expected retinal cell type markers expressed in each isolated population. (**e**) Images of dissociated mouse retinal cells after the SABER FISH protocol on dissociated mouse retinal cells. (**f**) A heatmap representing relative expression levels of BC subtype markers previously identified by scRNA sequencing that are differentially expressed (adjusted *p*-value*<*0.05) between *Grik1*^−^ and *Grik1*^+^ populations. (**g**) A representative scatter plot of log_2_ normalized counts of all genes (left panel) or BC2-BC4 marker genes (right panel) between Live *Grik1*^+^ cells and Probe-Seq *Grik1*^+^ Cells. Red line indicates a slope of 1. Select cell class-specific markers (Opn1mw, Apoe, and Rho) are labeled in red. (**h**) A heatmap of the stain index with varying levels of transcript expression and number of tiling oligonucleotides. The white spaces indicate a cutoff of SI*<*2. (**i**) Schematic and flow cytometry plots of iterative Probe-Seq. Three probe sets (*Gad1*, *Vsx2*, and *Grm6*) were hybridized to dissociated mouse retinal cells, and fluorescent oligonucleotides were hybridized only to *Gad1* and *Vsx2* probe sets to detect subsets of ACs (*Gad1*^+^; 3.54%) and BC/MG (*Vsx2*^+^; 11.9%). The fluorescent oligonucleotides were subsequently stripped with 50% formamide, which abolished the staining based on flow cytometry. Fluorescent oligonucleotides for *Grm6* were then hybridized to label a subset of BCs (*Grm6*^+^; 6.78%). HC, Horizontal Cell; RGC, Retinal Ganglion Cell; AC, Amacrine Cell; BC, Bipolar Cell; MG, Mller Glia; ONL, Outer Nuclear Layer; INL, Inner Nuclear Layer; GCL, Ganglion Cell Layer. Scale bars: 10 *µ*M (b, e).

To determine whether the isolated populations corresponded to the expected cell types, we reversed the crosslinking and extracted the RNA from these cells. SMART-Seq v.4 cDNA libraries were generated and sequenced on NextSeq 500. Each sample was sequenced to a mean of 15*±*3 million 75 bp paired-end reads to be able to reliably detect low abundance transcripts. The average 3’ bias for the mapped reads for all samples was 0.74 ± 0.02, which corresponds to a RNA Integrity Number (RIN) of 2-4^29^, indicating mild degradation of RNA. Unbiased hierarchical clustering showed that samples of the same cell population clustered together (average Pearson correlation between samples within population: r = 0.93) **(Supplementary Figure 1)**. The three populations were then analyzed for differential expression (DE). Between each population, the frequency distribution of all *p*-values showed an even distribution of null *p*-values, thus allowing for calculation of adjusted *p*-value using the Benjamini-Hochberg procedure **(Supplementary Figure 1)**. Between *Grik1*^−^ and *Grik1*^+^ populations, we found 1,740 differentially expressed genes (adjusted *p*-value *<* 0.05) out of 17,649 genes **(Supplementary Figure 1)**. The high number of genes detected indicates successful bulk RNA sequencing of low abundance transcripts.

To determine which retinal cell types were enriched in the isolated populations, we cross-referenced the DE gene set (adjusted *p*-value *<* 0.05) to retinal cell class-specific markers identified by Drop-Seq (see Methods for details of gene set curation)^3^. We saw that the *Vsx2*^−^ population was enriched for markers of all cell classes except for BCs and MG **(Supplementary Figure 2)**, as expected from the expression pattern of *Vsx2*^5^. The *Grik1*^−^ population was enriched for most BC and MG markers, while the *Grik1*^+^ population was enriched for a subset of BC markers **(Supplementary Figure 2)**. Accordingly, Gene Set Enrichment Analysis (GSEA) between *Vsx2*^−^ and *Grik1*^−^ populations indicated significant enrichment of rod, cone, AC, HC, and RGC markers in the *Vsx2*^−^ populations and BC and MG markers in the *Grik1*^−^ population (default significance at FDR *<* 0.25; Enrichment in *Vsx2*^−^ population: Rod: FDR *<* 0.001; Cone: FDR *<* 0.001; AC: FDR *<* 0.001; HC: FDR = 0.174; RGC: FDR = 0.224; Enrichment in *Grik1*^−^ population: BC: FDR *<* 0.001; MG: FDR *<* 0.001).

To determine which BC subtypes were enriched in the *Grik1*^−^ and *Grik1*^+^ populations, we cross-referenced the DE gene set (adjusted *p*-value *<* 0.05) to BC subtype specific markers identified by scRNA sequencing^5^. We found that the majority of BC2, BC3A, BC3B, and BC4 markers were enriched in the *Grik1*^+^ population as expected, and all other BC subtype markers were highly expressed in the *Grik1*^−^ population (Figure 1f). GSEA between *Grik1*^−^ and *Grik1*^+^ populations confirmed these results (Enrichment in *Grik1*^+^ population: BC2: FDR *<* 0.001; BC3A: FDR = 0.005; BC3B: FDR *<* 0.001; BC4: FDR *<* 0.001; Enrichment in *Grik1*^−^ population: BC1B: FDR = 0.132; BC5A: FDR = 0.135; BC5C: FDR = 0.136; BC5D: FDR = 0.172; BC6: FDR = 0.145; BC7: FDR = 0.169; BC8/9: FDR = 0.174; RBC: FDR *<* 0.001). From the DE analysis, we also identified the top 20 most DE genes that were specific to a cell population **(Supplementary Figure 3)**. We confirmed the expression of *Tpbgl*, a previously uncharacterized transcript, in *Grik1*^+^ cells by SABER FISH in retinal tissue sections **(Supplementary Figure 3)**. These results indicate that the cell populations isolated and profiled by Probe-Seq correspond to the expected BC subtypes.

We next aimed to determine the relative quality of the transcriptomes obtained by Probe-Seq versus those obtained from freshly dissociated cells. To this end, we electroporated the Grik1^CRM4^-GFP reporter plasmid into the developing retina at P2. We previously showed that 72% of GFP^+^ cells were *Grik1*^+^ using this reporter^24^. At P40, the retinas were harvested, and the electroporated region was dissociated into a single cell suspension. GFP^+^ cells were FACS isolated into Trizol (henceforth called Live cells). Simultaneously, cells from the unelectroporated region from the same retina were used for Probe-Seq of *Grik1*^+^ cells (henceforth called Probe-Seq cells). On average, 836±155 Live cells (n=3) and 10,000±0 Probe-Seq cells (n=3), were collected for transcriptional analysis. A strong correlation of expression levels of all genes between Live and Probe-Seq cells was seen (average Pearson correlation r = 0.78±0.01) (Figure 1g). Since 72% of the GFP^+^ population in the Live cell population were expected to be *Grik1*^+^ cells, we expected that the correlation for markers of BC2-BC4 to be approximately 0.72. Notably, when analyzing only the subtype-specific markers, the correlation was 0.66 between Live and Probe-Seq cells, with enrichment of these markers in the Probe-Seq population (Figure 1g). Therefore, expression of GFP in *Grik1*^−^ cells using the Grik1^CRM4^-GFP reporter plasmid could explain, at least partially, the discrepancy between the Probe-Seq and Live cell populations. Accordingly, DE analysis between these two populations indicated enrichment of rod, cone, and MG markers, as evidenced by high expression of marker genes (i.e. Rho, Opn1mw, and Apoe) in the Live cell population (Figure 1g). Despite the imperfect match in the cellular compositions of these two preparations, the strong correlation between the transcriptomes obtained by Probe-Seq and traditional live cell sorting indicates that high quality transcriptomes can be obtained by this method.

We next sought to investigate the parameters for SABER probe sets that are important for successful FACS isolation. We reasoned that the ability to resolve a targeted cell from the total cell population would be dependent upon the total number of fluorescent probes in that cell, which can be increased by targeting more tiling oligonucleotides to each transcript, or by targeting more abundant RNA species. To investigate these parameters, we generated three gene-specific probe sets, for *Grik1*, *Grm6*, and *Neto1*, which exhibit high-to-low levels of gene expression based upon FISH analysis in retina tissue sections (Number of puncta per positive cell: *Grik1*: 37; *Grm6*: 20; *Neto1*: 6.5) **(Supplementary Figure 4)**. For each gene-specific probe set, 48, 24, or 12 randomly-chosen tiling oligonucleotides were pooled for extension. These were then used for Probe-Seq, and the fluorescent signals from the FACS were analyzed to calculate the Stain Index (SI; see Methods for calculation of SI). This allowed for quantification of the separation of the positive population from the negative population. The SI was found to decrease with the reduced number of tiling oligonucleotides and the level of expression of each gene (Figure 1h). However, with an SI cutoff of 2, 12 tiling oligonucleotides were sufficient for confidence in the separation of gene-positive and negative populations, demonstrating that short transcripts or few tiling oligonucleotides can be used successfully for Probe-Seq.

To label multiple cell types, or cellular states, it is often necessary to use a combination of gene-specific probe sets. Serial detection of markers has been achieved using SABER-FISH (Exchange-SABER). Exchange-SABER has enabled the labeling of seven retinal cell classes using three cycles of FISH. After each round of imaging, the fluorescent oligonucleotides are stripped using conditions that do not strip the gene-specific probe sets, allowing fluorescent channels to be reused for detection of different genes in the same cells. To determine whether serial multiplexing is feasible with Probe-Seq, dissociated mouse retinal cells were incubated with three gene-specific probe sets for *Gad1*, *Vsx2*, and *Grm6* (Figure 1i). For the first round of flow cytometry, the fluorescent oligonucleotides for detecting *Gad1* and *Vsx2* were applied. These were assayed by flow cytometry and then stripped using 50% formamide. The removal of the fluorescent oligonucleotides was confirmed by the lack of signal in a subsequent round of flow cytometry (Figure 1i). We then applied the fluorescent oligonucleotides for *Grm6* and were able to detect a new population of cells from the same cellular pool. These results indicate that serial multiplexed Probe-Seq can allow detection of multiple cell types in the same cell preparation with iterative rounds of hybridization and FACS.

### Probe-Seq enables isolation and RNA sequencing of cell type-specific nuclei from frozen postmortem human tissue

To determine whether Probe-Seq will allow one to access the transcriptomes of the many archived human tissue samples, we tested the method on frozen human retinas. Nuclear preparations were made, as whole cell approaches to frozen cells are not feasible^30,31^. The initial test was carried out on frozen mouse retinas. Nuclei were extracted by Dounce homogenization, fixed with 4% PFA, and labeled by a gene-specific probe set for *Grik1* **(Supplementary Figure 5)**. *Grik1*^HI^ (not *Grik1*^MID^) and *Grik1*^−^ populations were isolated by FACS, the nuclear RNA was extracted, and the cDNA was sequenced. We cross-referenced the DE gene set (adjusted *p*-value *<* 0.05) to BC subtype specific markers and found that the majority of mouse BC3A, BC3B, and BC4 markers were enriched in the *Grik1*^+^ population, as expected, and all other BC subtype markers were highly expressed in the *Grik1*^−^ population **(Supplementary Figure 5)**. These results indicate that cell type-specific nuclear RNA from frozen tissue can be isolated by Probe-Seq.

We thus obtained fresh-frozen human retinas (age range: 40 60; see Methods for full description of samples), and aimed to isolate and profile human BC subtypes using a probe set for *GRM6*, which is expressed in cone ON bipolar cells and rod bipolar cells (RBC) in the mouse and human retina^5,32^ (Figure 2a). To test the *GRM6* probe set in human retinas, it was first applied to a fixed human tissue section, where signal was observed in the expected pattern, in a subset of cells in the inner nuclear layer, where BCs reside (Figure 2b). Nuclei were extracted from frozen human peripheral retinas, fixed, and incubated with the *GRM6* probe set. The *GRM6*^−^ and *GRM6*^+^ nuclei were then isolated by FACS after application of the fluorescent oligonucleotides (Figure 2c). On average, 43,000±35,500 *GRM6*^−^ nuclei and 1,800±781 *GRM6*^+^ nuclei were isolated from approximately 5 mm x 5 mm square of the retina per biological replicate (Figure 2d). SMART-Seq v.4 cDNA libraries were sequenced on NextSeq 500, with each sample sequenced to a mean depth of 18±3 million 75 bp paired-end reads. The average 3’ bias for the mapped reads of the negative population was 0.70±0.04, indicating slight degradation of RNA. Quality control of the read mapping and DE analysis indicated successful RNA sequencing and DE analysis **(Supplementary Figure 6)**. Upon filtering out genes with zero counts in more than 4 samples, 1,956 out of 9,619 genes were differentially expressed (adjusted *p*-value *<* 0.05).

**Figure 2:**
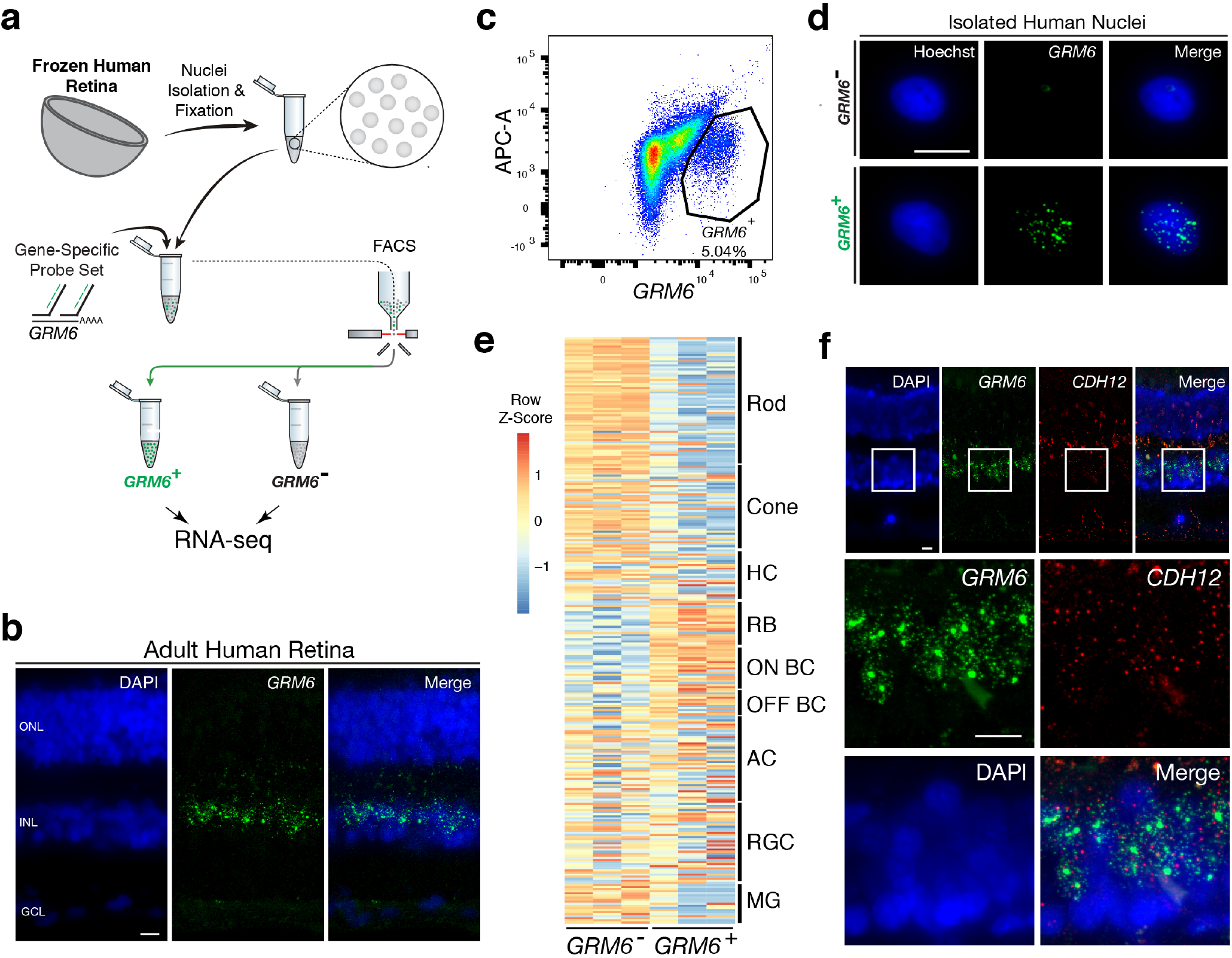
Transcriptional profiling of nuclear RNA isolated from specific BC subtypes from frozen human retina. (**a**) Schematic of Probe-Seq for the fresh frozen adult human retina. Single nuclei were prepared and then fixed. Nuclei were incubated with a SABER probe set for *GRM6* and then incubated with fluorescent oligonucleotides. *GRM6*^+^ and *GRM6*^−^ populations were isolated by FACS for downstream RNA sequencing. (**b**) Image of an adult human retina section probed with a SABER *GRM6* probe set. (**c**) FACS plot of all single nuclei with *GRM6* fluorescence on the x-axis and empty APC-A autofluorescence on the y-axis. (**d**) Images of isolated nuclei processed using SABER FISH for *GRM6*. (**e**) A heatmap representing relative expression levels of human retinal cell type markers previously identified by scRNA sequencing that are differentially expressed (adjusted *p*-value*<*0.05) between *GRM6*^−^, and *GRM6*^+^ populations. (**f**) High and low magnification images of a human retinal section after the SABER FISH protocol for *CDH12*, a highly-enriched transcript in the *GRM6*^+^ population. HC, Horizontal Cell; RGC, Retinal Ganglion Cell; AC, Amacrine Cell; ON BC, ON Bipolar Cell; RBC, Rod Bipolar Cell; OFF BC, OFF Bipolar Cell; MG, Mller Glia; ONL, Outer Nuclear Layer; INL, Inner Nuclear Layer; GCL, Ganglion Cell Layer. Scale bars: 10 *µ*M (b, d, f).

We compared the DE gene set (adjusted *p*-value *<* 0.05) to human retinal cell type-specific markers identified by scRNA sequencing^32^. We found an enrichment of markers for RBCs and ON BCs in the *GRM6*^+^ population (Figure 2e). GSEA between *GRM6*^−^ and *GRM6*^+^ confirmed these results (Enrichment in *GRM6*^+^ population: RBC: FDR = 0.003; ON BC-1: FDR = 0.181; ON BC-2: FDR = 0.126), indicating that the expected human retinal populations were accurately isolated. We validated the expression of *CDH12*, a transcript highly enriched in the *GRM6*^+^ population, but previously not reported to be a marker of these subtypes, by performing SABER FISH on fixed adult human retina sections (Figure 2f). These results show that nuclear transcripts isolated from frozen tissue by Probe-Seq are sufficient for transcriptional profiling.

### Isolation and transcriptional profiling of intestinal stem cells from the *Drosophila* midgut

To determine whether Probe-Seq can be successfully applied to non-CNS cells, and to cells from invertebrates, we applied the method to the midgut of *Drosophila melanogaster*. The adult *Drosophila* midgut is composed of four major cell types - enterocytes (EC), enteroendocrine cells (EE), enteroblasts (EB), and intestinal stem cells (ISC), though recent profiling studies have revealed heterogeneity among ECs and ISC/EBs^33,34^. We aimed to isolate ISCs and EBs using a gene-specific probe set for escargot (*esg*), a well-characterized marker for these cell types (Figure 3a). As SABER-FISH had not yet been tested on *Drosophila* tissue, we first tested this method on wholemounts of the *Drosophila* gut. SABER FISH signal was observed in the appropriate pattern, in a subset of midgut cells (Figure 3b). To perform Probe-Seq, we dissociated 35-40 *Drosophila* midguts per biological replicate, fixed in 4% PFA, and incubated the cells with the *esg* probe set. FACS was then used to sort *esg*^−^ and *esg*^+^ populations (Figure 3c-d). As only the 2N single cells were sorted, proliferating ISCs and polyploid ECs were excluded. On average, 1,400±208 *esg*^−^ cells and 1,000±404 *esg*^+^ cells per biological replicate were isolated. SMART-Seq v.4 cDNA libraries were sequenced on NextSeq 500 to a mean of 16±4 million 75 bp paired-end reads. Quality control of the read mapping and DE analysis indicated successful RNA sequencing and DE analysis **(Supplementary Figure 7)**. Upon filtering out genes with zero counts in more than 3 samples, 405 out of 1,596 genes were differentially expressed (adjusted *p*-value *<* 0.05).

**Figure 3:**
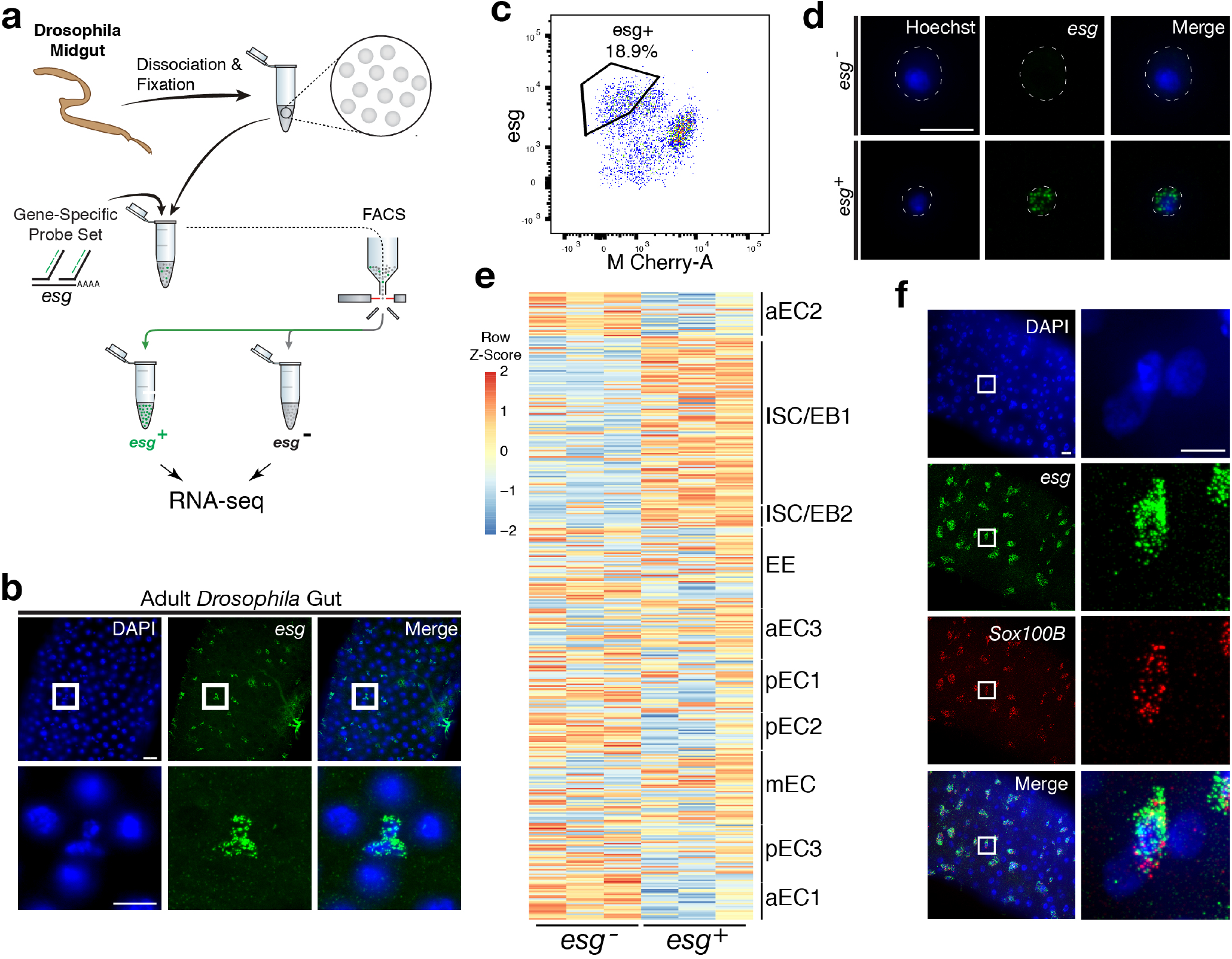
Isolation and transcriptional profiling of ISC/EBs from the adult Drosophila midgut. (**a**) Schematic of Probe-Seq for the adult Drosophila midgut. Midguts from 7-10-day old female flies were dissociated into single cells and fixed. Cells were incubated with a SABER FISH probe set for *esg* and subsequently incubated with fluorescent oligonucleotides. *esg*^+^ and *esg*^−^ populations were isolated by FACS for downstream RNA sequencing. (**b**) Image of a wholemount adult Drosophila midgut following the SABER FISH protocol using an *esg* probe set. (**c**) FACS plot of all single cells with *esg* fluorescence on the y-axis and empty M-Cherry-A autofluorescence on the x-axis. (**d**) Images of isolated midgut cells processed using SABER FISH for *esg*. The white dotted lines demarcate cell outlines. (**e**) A heatmap representing relative expression levels of differentially expressed (adjusted *p*-value*<*0.05) genes for *Drosophila* gut cell type markers previously identified by scRNA sequencing, between *esg*^−^, and *esg*^+^ populations. (**f**) Images of a *Drosophila* midgut wholemount after the SABER FISH protocol for an ISC/EB marker, *Sox100B*, a highly-enriched transcript in the *esg*^+^ population. EC, Enterocyte; ISC, Intestinal Stem Cell; EB, Enteroblast; EE, Enteroendocrine Cell. Scale bars: 10 *µ*M (d, f, right panels); 20 *µ*M (b, upper panels f, left panels).

The DE gene set (adjusted *p*-value *<* 0.05) from Probe-Seq was compared to cell type-specific markers identified by scRNA sequencing^34^. An enrichment of ISC/EB1 and ISC/EB2 markers was observed in the *esg*^+^ population isolated using Probe-Seq, while markers of all other cell types were enriched in the *esg*^−^ population (Figure 3e). GSEA between *esg*^+^ and *esg*^−^ populations isolated using Probe-Seq indicated significant enrichment of ISC/EB1 and ISC/EB2 markers in the *esg*^+^ population and all other cell type markers in the *esg*^−^ population (Enrichment in *esg*^+^ population: ISC/EB1: FDR *<* 0.001; ISC/EB2: FDR = 0.081; Enrichment in *esg*^−^ population: aEC1: FDR *<* 0.001; aEC2: FDR *<* 0.001; pEC2 FDR *<* 0.001; pEC1: FDR *<* 0.001; pEC3: FDR *<* 0.001; EE: FDR *<* 0.001; aEC3: FDR = 0.009; mEC: FDR = 0.041). Using SABER FISH on wholemounts of midguts, we validated the co-localization of *esg* and *Sox100B*, a transcript significantly enriched in the *esg*^+^ population (Figure 3f). Additionally, we cross-referenced the Probe-Seq DE gene set (adjusted *p*-value *<* 0.05) to ISC/EB and EC markers defined by DamID profiling of the adult *Drosophila* gut^35^. From this analysis, the majority of ISC/EB and EC markers were seen to be enriched in *esg*^+^ and *esg*^−^ populations, respectively **(Supplementary Figure 8)**. These results demonstrate that Probe-Seq enables the isolation and transcriptional profiling of specific cell types from invertebrate non-CNS tissue.

### Transcriptome profiling of the central chick retina reveals unique transcripts expressed in cells that give rise to the high acuity area

The central chicken retina contains a region thought to endow high acuity vision, given its cellular composition and arrangement of cells. It comprises a small and discrete area that is devoid of rod photoreceptors and enriched in cone photoreceptors, with a high density of retinal ganglion cells, the output neurons of the retina^36,37^. These features are shared with the high acuity areas (HAA) of other species, including human. Although we have shown that *FGF8*, *CYP26C1*, and *CYP26A1* are highly enriched in this area at embryonic day 6 (E6)^37^, the other molecular determinants that may play a role in HAA development are unknown. Probe-Seq was thus used to isolate and sequence *FGF8*^+^ cells. Hamburger-Hamilton stage 28 (HH28) chick retinas were dissociated into single cells, fixed, and probed for the *FGF8* transcript (Figure 4a). On average, 7,000±6,950 *FGF8*^+^ cells and 189,000±19,000 *FGF8*^−^ cells were FACS isolated into individual populations (Figure 4b). The cDNA from each population (n=3) was sequenced to a mean depth of 18*±*7 million 75bp paired-end reads on NextSeq 500. Interestingly, the 3’ bias of the mapped reads from the chick retina was significantly reduced compared to that of the mouse retina (0.59±0.04, comparable to RIN 6-8) **(Supplementary Figure 9)**. Quality control of the read mapping and DE analysis indicated successful RNA sequencing and DE analysis **(Supplementary Figure 10)**. Between *FGF8*^−^ and *FGF8*^+^ populations, we found 1,924 DE genes out of 12,053 genes.

**Figure 4:**
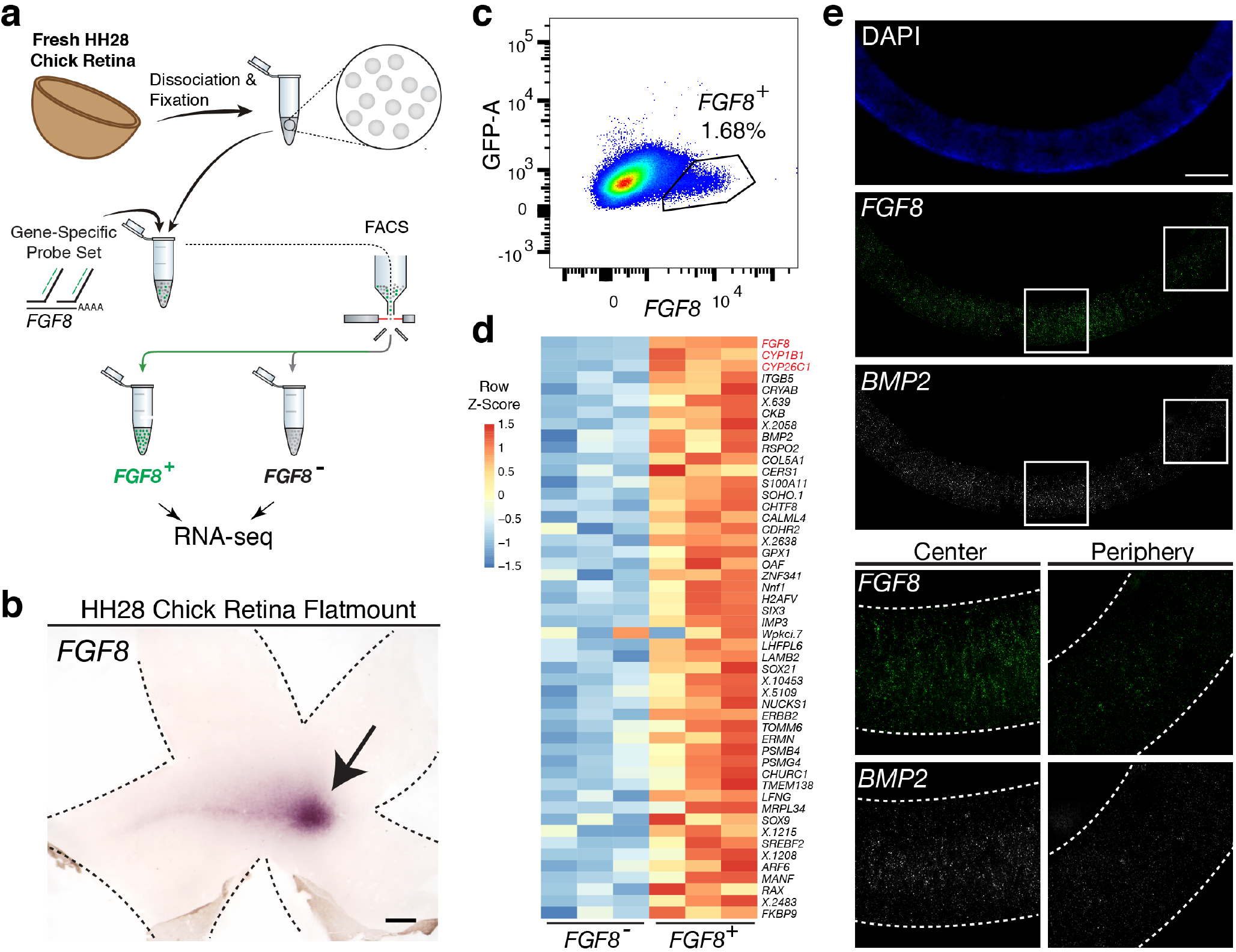
Probe-Seq identifies the transcriptional landscape of chick central progenitor cells that give rise to the high acuity area. (**a**) Schematic of Probe-Seq for the developing HH28 chick retina. The chick retina was dissociated into single cells and fixed. Cells were incubated with a SABER FISH probe set for *FGF8* and subsequently incubated with fluorescent oligonucleotides. *FGF8*^+^ and *FGF8*^−^ populations were isolated by FACS for downstream RNA sequencing. (**b**) FACS plot of all single cells with *FGF8* fluorescence on the x-axis and empty GFP-A autofluorescence on the y-axis. (**c**) A heatmap of unbiased top 50 genes that were enriched in the *FGF8*^+^ population compared to the *FGF8*^−^ population. (**d**) Images of a section spanning the central HH28 chick retina after the SABER FISH protocol for *FGF8* and *BMP2*, a transcript highly enriched in the *FGF8*^+^ population, Scale bars: 500 *µ*M (b); 50 *µ*M (e).

Among the top 50 most enriched DE transcripts in the *FGF8*^+^ population were *FGF8*, *CYP1B1*, and *CYP26C1* (Figure 4c). The latter two transcripts are components of the retinoic acid signaling pathway, and were previously shown to be highly enriched in the central retina where *FGF8* is expressed^37^. Previously, *FGF8* expression was shown to be largely confined to the area where progenitor cells reside. However, it was unclear whether it was also expressed in differentiated cells^37^. DE analysis of the *FGF8*^−^ and *FGF8*^+^ populations revealed enrichment of early differentiation markers of RGCs (i.e. *NEFL*) and photoreceptors (i.e. *NEUROD1*) in the *FGF8*^−^ population, confirming that *FGF8* is mostly expressed in central progenitor cells **(Supplementary Figure 11)**. As we wished to validate the DE genes using FISH on sections, and SABER FISH had not yet been tested on chick tissue, we first tested the method using the *FGF8* probe set on chick tissue sections (Figure 4e). Robust and specific FISH signal was seen in the appropriate pattern for *FGF8*. An additional SABER-FISH probe set for *BMP2*, a transcript enriched in the *FGF8*^+^ population, was then used on developing HH28 chick central retinal sections. *BMP2* was found to be highly enriched within the *FGF8*^+^ population, i.e expression was confined to a discrete central retina where *FGF8* was expressed (Figure 4e). These results indicate that Probe-Seq of the developing chick retina using *FGF8* as a molecular handle can reveal the transcriptional profile of the progenitor cells that will comprise the chick high acuity area.

## DISCUSSION

Studies of model organisms have allowed the dissection of molecular mechanisms that underlie a variety of biological processes. However, each organism across the evolutionary tree possesses unique traits, and understanding these traits will greatly enrich our understanding of biological processes. Non-model organisms can now be investigated at the genetic level, due to advances in DNA sequencing, transcriptional profiling, and genome modification methods. Despite progress, challenges remain to achieve greater depth in the characterization of the transcriptomes of rarer cell types within heterogeneous tissues. Even in model organisms, deep transcriptional profiling of specific cell types remains difficult if specific cis-regulatory elements are unavailable. To overcome these challenges, we developed Probe-Seq.

Probe-Seq uses a FISH method based upon SABER probes to hybridize gene-specific probe sets to RNAs of interest^24^. This method provides amplified fluorescent detection of RNA molecules and can be spectrally or serially multiplexed to mark cell populations based on combinatorial RNA expression profiles. Previously-identified markers can thus be targeted by gene-specific probe sets to isolate specific cell types by FACS. Subsequently, deep RNA sequencing can be carried out on the sorted population to generate cell type-specific transcriptome profiles. Due to the reliance on RNA for cell sorting, rather than protein, this method is applicable across organisms. Other RNA-based methods to label specific cell types for downstream sequencing have not been tested with tissue samples, and/or require cell encapsulation in a microfluidic device^25,38,39^.

Probe-Seq allowed the isolation and profiling of RNA from fresh mouse, frozen human, and fresh chick retinas, as well as gut cells from *Drosophila melanogaster*. Aside from the different dissociation protocols, Probe-Seq does not require species- or tissue-specific alterations. To profile multiple cellular subtypes, serial multiplexed Probe-Seq allows for iterative labeling, sorting, and re-labeling. This strategy enables separation of FACS-isolated, broad populations into finer sub-populations. The Probe-Seq method is also cost and time effective, with less than 6 hours of hands-on time, including FISH, FACS, and library preparation. Per sample, we estimate the cost to be less than $200, from start to finish, achieving 15 million paired-end reads. The Probe-Seq protocol may be further optimized to maximize utility. For example, the protocol may be further modified to use other single molecule FISH methods such as clampFISH^40^ or RNAscope^41^ rather than SABER-FISH, as these methods have their respective strengths and weaknesses. Additionally, the protocol may also be adapted to use a cell strainer for the wash steps to minimize cell loss from centrifugation. Further development for scRNA sequencing after cell type enrichment using Probe-Seq may also be possible. For this, however, adaptation of Probe-Seq for the reversal of crosslinks for scRNA sequencing^42–44^ will likely be necessary.

## METHODS

### Mouse retina samples

All animals were handled according to protocols approved by the Institutional Animal Care and Use Committee (IACUC) of Harvard University. For fresh samples, retinas of adult CD1 mice (*>*P30) from Charles River Laboratories were dissected. For frozen samples, retinas of adult CD1 mice (*>*P30) were dissected and frozen in a slurry of isopentane and dry ice and kept at −80°C.

### Human retina samples

Frozen eyes were obtained from Restore Life USA (Elizabethton, TN) through TissueForResearch. Patient DRLU041818C was a 53-year-old female with no clinical eye diagnosis and the postmortem interval was 9 hours. Patient DRLU051918A was a 43-year-old female with no clinical eye diagnosis and the postmortem interval was 5 hours. Patient DRLU031318A was a 47-year-old female with no clinical eye diagnosis and the postmortem interval was 7 hours. This IRB protocol (IRB17-1781) was determined to be not-human subject research by the Harvard University-Area Committee on the Use of Human Subjects.

### Chick retina samples

Fertilized White Leghorn eggs from Charles River Laboratories were incubated at 38°C with 40% humidity. Embryos were staged according to Hamburger and Hamilton up to HH28^45^.

### *Drosophila melanogaster* midgut samples

Tissues were dissected from female 7-10-day-old adult Oregon-R *Drosophila melanogaster*. Flies were reared on standard cornmeal/agar medium in 12:12 light:dark cycles at 25°C.

### Dissociation of mouse and chick retinas

Mouse or chick retinas were dissected away from other ocular tissues in Hanks Balanced Salt Solution (Thermo Fisher Scientific, cat. #14025092) or PBS. The retina was then transferred to a microcentrifuge tube and incubated for 7 minutes at 37°C with an activated papain dissociation solution (87.5 mM HEPES pH 7.0 (Thermo Fisher Scientific, cat. #15630080), 2.5 mM L-Cysteine (MilliporeSigma, cat. #168149), 0.5 mM EDTA pH 8.0 (Thermo Fisher Scientific, cat. #AM9260G), 10 *µ*L Papain Suspension (Worthington, cat. #LS0003126), 19.6 *µ*L UltraPure Nuclease-Free Water (Thermo Fisher Scientific, cat. #10977023), HBSS up to 400 *µ*L, activated by a 15-minute incubation at 37°C). The retina was then centrifuged at 600 xg for 3 minutes. The supernatant was removed, and 1 mL of HBSS/10% FBS (Thermo Fisher Scientific, cat. #10437028) was added without agitation to the pellet. The pellet was centrifuged at 600 xg for 3 minutes. The supernatant was removed, and 600 *µ*L of trituration buffer (DMEM (Thermo Fisher Scientific, cat. #11995065), 0.4% (wt/vol) Bovine Serum Albumin (MilliporeSigma cat. #A9418)) was added. The pellet was dissociated by trituration at room temperature (RT) using a P1000 pipette up to 20 times or until the solution was homogenous.

### Dissociation of *Drosophila* midgut

35-40 *Drosophila* midguts were dissected in PBS and transferred to 1% BSA/PBS solution. The midguts were incubated in 400 *µ*L of Elastase/PBS solution (1 mg/mL, MilliporeSigma cat. #E0258) for 30 minutes to 1 hour at RT, with trituration with a P1000 pipette every 15 minutes. 1 mL of 1% BSA/PBS was then added. This solution was overlaid on top of Optiprep/PBS (MilliporeSigma, cat. #D1556) solution with a density of 1.12 g/mL in a 5-mL polypropylene tube (Thermo Fisher Scientific, cat. #1495911A). The solution was centrifuged at 800 xg at RT for 20 minutes. The top layer with viable cells was collected for further processing.

### Mouse and human frozen nuclei isolation

Upon thawing, tissue was immediately incubated in 1% PFA (with 1 *µ*L mL^−1^ RNasin Plus (Promega, cat. #N2611)) for 5 minutes at 4°C. Nuclei were prepared by Dounce homogenizing in Homogenization Buffer (250 mM sucrose, 25 mM KCl, 5 mM MgCl_2_, 10mM Tris buffer, pH 8.0, 1 *µ*M DTT, 1x Protease Inhibitor (Promega, cat. #6521), Hoechst 33342 10 ng mL^−1^ (Thermo Fisher Scientific, cat. #H3570), 0.1% Triton X-100, 1 *µ*L mL^−1^ RNasin Plus). Sample was then overlaid on top of 20% sucrose solution (25 mM KCl, 5 mM MgCl_2_, 10mM Tris buffer, pH 8.0) and spun at 500 xg for 12 minutes at 4°C.

### Probe-Seq for whole cells and nuclei

For all solutions, 1 *µ*L mL^−1^ RNasin Plus was added 10 minutes before use. If the cells or nuclei were not already pelleted, the suspended cells/nuclei (henceforth called cells) were centrifuged at 600 xg for 5 minutes at 4°C. The cells were then resuspended in 1 mL of 4% PFA (Electron Microscopy Sciences, cat. #15714S, diluted in PBS) and incubated at 4°C for 15 minutes with rocking. The cells were centrifuged at 2000 xg for 5 minutes at 4°C. Except in the case of nuclei, the supernatant was removed, and the cells were resuspended in 1 mL of Permeabilization Buffer (Hoechst 33342 10 *µ*g mL^−1^, 0.1% Trixon X-100, PBS up to 1 mL) and incubated for 10 minutes at 4°C with rocking. For both cells and nuclei, the cells were next centrifuged at 2000 xg for 5 minutes at 4°C. The supernatant was removed, and the cells were resuspended in 500 *µ*L of pre-warmed (43°C) 40% wash Hybridization solution (wHyb; 2x SSC (Thermo Fisher Scientific, cat. #15557044), 40% deionized formamide (MilliporeSigma, cat. #S4117), diluted in UltraPure Water). Compared to the original SABER protocol^24^, the Tween-20 was removed as we found that it causes cell clumping. The cells were incubated for at least 30 minutes at 43°C. After this step, the cell pellet became transparent. The cells were centrifuged at 2000 xg for 5 minutes at RT, and the supernatant was carefully removed, leaving ~100 *µ*L of supernatant. The cells were then resuspended in 100 *µ*L of pre-warmed (43°C) Probe Mix (1 *µ*g of probe per gene, 96 *µ*L of Hyb1 solution (2.5x SSC, 50% deionized formamide, 12.5% Dextran Sulfate (MilliporeSigma cat. #D8906)), diluted up to 120 *µ*L with UltraPure Water) and incubated overnight (16-24 hours) at 43°C.

500 *µ*L of pre-warmed (43°C) 40% wHyb was added to the cells and centrifuged at 2000 xg for 5 minutes at RT. The supernatant was removed, and the cells were resuspended in 500 *µ*L of pre-warmed (43°C) 40% wHyb. The cells were incubated for 15 minutes at 43°C. The cells were then centrifuged at 2000 xg for 5 minutes at RT, and the supernatant was removed. 1 mL of pre-warmed (43°C) 2x SSC solution was added, the cells were resuspended, and incubated for 5 minutes at 43°C. The cells were centrifuged at 2000 xg for 5 minutes at RT, and the supernatant was removed. Cells were then resuspended in 500 *µ*L of pre-warmed (37°C) PBS. The cells were centrifuged at 2000 xg for 5 minutes at RT, and the supernatant was removed. The cells were resuspended in 100 *µ*L of Fluorescent Oligonucleotide Mix (100 *µ*L of PBS, 2 *µ*L of each 10 *µ*M Fluorescent Oligonucleotide) and incubated for 10 minutes at 37°C. After incubation, 500 *µ*L of pre-warmed (37°C) PBS was added and the cells were centrifuged at 2000 xg for 5 minutes at RT. The supernatant was removed, and the cells were resuspended in 500 *µ*L of pre-warmed (37°C) PBS. The cells were incubated for 5 minutes at 37°C. The cells were centrifuged at 2000 xg for 5 minutes at RT, the supernatant was removed, and the cells were resuspended in 500-1000 *µ*L of PBS, depending on cell concentration.

### FACS isolation of specific cell types

The suspended labeled cells were kept on ice before FACS. Immediately before FACS, the cells were filtered through a 35 *µ*M filter (Thermo Fisher Scientific, cat. #352235) for mouse, chick, and human retina cells/nuclei or a 70 *µ*M filter (Thermo Fisher Scientific, cat. #352350) for *Drosophila* cells. FACSAria (BD Biosciences) with 488, 561, 594, and 633 lasers was used for the sorts. 2N single cells were gated based on the Hoechst histogram. For the *Drosophila* gut, debris was gated out first by FSC/SSC plot because the high number of debris events masked the Hoechst^+^ peaks. Out of the single cells, a 2-dimensional plot (with one axis being the fluorescent channel of interest and another axis that is empty) was used to plot the negative and positive populations. The events that ran along the diagonal in this plot were considered negative, and the positive events were either left- or right-shifted (depending on axes) compared to the diagonal events. For some samples, the number of sorted cells was capped, as indicated by a standard deviation of 0. Different populations were sorted into microcentrifuge tubes with 500 *µ*L of PBS and kept on ice after FACS. The protocol was later modified so that the cells are sorted into 500 *µ*L of 1% BSA/PBS, as this significantly improved cell pelleting. The data obtained for this study did not use 1% BSA/PBS.

### RNA isolation and library preparation

The sorted cells were transferred to a 5-mL polypropylene tube and centrifuged at 3000 xg for 7 minutes at RT. The supernatant was removed as much as possible, the cells were resuspended in 100 *µ*L Digestion Mix (RecoverAll Total Nuclear Isolation Kit (Thermo Fisher Scientific, cat. #AM1975) 100 *µ*L of Digestion Buffer, 4 *µ*L of protease), and incubated for 3 hours at 50°C, which differs from the manufacturer’s protocol. The downstream steps were according to the manufacturer’s protocol. The volume of ethanol/additive mix in the kit was adjusted based on the total volume (100 *µ*L of Digestion Mix and remaining volume after cell pelleting). The libraries for RNA sequencing were generated using the SMART-Seq v.4 Ultra Low Input RNA kit (Takara Bio, cat. #634890) and Nextera XT DNA Library Prep Kit (Illumina, cat. #FC1311096) according to the manufacturer’s protocol. The number of cycles for SMART-Seq v.4 protocol was as follows: Mouse *Vsx2/Grik1*: 13 cycles; Chick *FGF8*: 16 cycles; Human *GRM6*: 16 cycles; *Drosophila esg*: 17 cycles. 150 pg of total cDNA was used as the input for Nextera XT after SMART-Seq v.4, and 12 cycles were used except for *Drosophila* samples for which 14 cycles were necessary. The cDNA library fragment size was determined by the BioAnalyzer 2100 HS DNA Assay (Agilent, cat. #50674626). The libraries were sequenced as 75bp paired-end reads on NextSeq 500 (Illumina).

### RNA-Seq data analysis

Quality control of RNA-seq reads were performed using fastqc version 0.10.1 (https://www.bioinformatics.babraham.ac.uk/projects/fastqc/). RNA-seq reads were clipped and mapped onto the either the mouse genome (Ensembl GRCm38.90), human genome (Ensembl GRCh38.94), chick genome (Ensembl GRCg6a.96), or *Drosophila* genome (BDGP6.22) using STAR version 2.5.2b^46,47^. Parameters used were as follows: –runThreadN 6 –readFilesCommand zcat –outSAMtype BAM SortedByCoordinate – outSAMunmapped Within –outSAMattributes Standard –clip3pAdapterSeq – quantMode TranscriptomeSAM GeneCounts

Alignment quality control was performed using Qualimap version 2.2.1^48^. Read counts were produced by HT-seq version 0.9.1^49^. Parameters used were as follows: -i gene name -s no

The resulting matrix of read counts were analyzed for differential expression by DESeq2 version 3.9^50^. For the DE analysis of human and *Drosophila* samples, any genes with more than 4 and 3 samples with zero reads, respectively, were discarded. The R scripts used for differential expression analysis are available in Supplementary Files.

### Gene set curation

Unique marker genes that define different cell types in different tissue types were curated in an unbiased manner. For the mouse retina, marker genes of major cell types were identified from scRNA sequencing^3^. Genes that were found in more than one cluster were removed from the analysis to obtain unique cluster-specific markers. Rod-specific genes were highly represented in all clusters; thus, they were considered non-unique by this analysis. Therefore, top 20 rod-specific genes were manually added after non-unique genes were removed. For mouse BCs, marker genes of BC subtypes with high confidence were identified by scRNA sequencing^5^. Genes that were found in more than one cluster was removed from the analysis. For the human retina, marker genes of major cell types were identified by scRNA sequencing^32^. Marker genes that were expressed in *<* 90% of cells in the cluster were removed for analysis. For the *Drosophila* gut, marker genes of major cell types were identified from DamID transcriptional profiling and scRNA sequencing^34,35^. For the DamID dataset, a cutoff of FDR *<* 0.01 was used for marker genes that were specifically expressed between ISC/EBs and ECs.

### Gene set enrichment analysis

GSEAPreranked analysis was performed using GSEA v3.0^51^. Curated gene sets described above were used to define various cell types. Parameters used were as follows: Number of permutations: 1000; Enrichment statistic: classic; the ranked file was generated using log_2_FoldChange generated by DESeq2. To determine significance, we used the default FDR *<* 0.25 for all gene sets.

### SABER probe synthesis

SABER probe sets were synthesized using the original protocol^24^. The gene of interest was searched in the UCSC Genome Browser. Then, the BED files for genes of interest were generated through the UCSC Table Browser with the following parameters: group: Genes and Gene Predictions; track: NCBI RefSeq; table: UCSC RefSeq (refGene); region: position; output format: BED; Create one BED record per: Exons. If multiple isoforms were present in the BED output file, all but one was removed manually. Genome-wide probe sets for mouse, human, chick, and *Drosophila* were downloaded from (https://oligopaints.hms.harvard.edu/genome-files) with Balance setting. The tiling oligonucleotide sequences were generated using intersectBed (bedtools 2.27.1) between the BED output file and the genome-wide chromosome-specific BED file with f 1. If the BED file sequences were on the + strand, the reverse complement probe set was generated using OligoMiners probeRC.py script (https://github.com/brianbeliveau/OligoMiner). For each tiling oligonucleotide sequence, hairpin primer sequences were added following a TTT linker. The tiling oligonucleotides were ordered from IDT with the following specifications in a 96-well format: 10 nmole, resuspended in IDTE pH 7.5, V-Bottom Plate, and normalized to 80 *µ*M. The tiling oligonucleotides were then combined into one pool for gene-specific probe set synthesis using a multi-channel pipette. The tiling oligonucleotide sequences for every gene-specific probe set used in this study are provided in Supplementary Files.

Tiling oligonucleotides were extended by a PER concatemerization reaction (1X PBS, 10 mM MgSO_4_, dNTP (0.3 mM of A, C, and T), 0.1 *µ*M Clean.G, 0.5 *µ*L Bst Polymerase (McLab, cat. #BPL-300), 0.5 *µ*M hairpin, 1 *µ*M oligonucleotide pool). The reaction was incubated without the tiling oligonucleotide pool for 15 minutes at 37°C. Then, the oligonucleotide pool was added and the reaction was incubated for 100 minutes at 37°C, 20 minutes at 80°C, and incubated at 4°C until probe set purification. 8 *µ*L of the reaction was analyzed on an 1.25% agarose gel (run time of 8 minutes at 150 volts) to confirm the probe set length. Probe sets of 300-700 nt were used for the study. The 37°C extension time was increased or decreased (from 100 minutes) based on desired concatemer length. The probe set was purified using MinElute PCR Purification Kit (Qiagen, cat. #28004) following manufacturers protocol. The probe set was eluted in 25 *µ*L UltraPure Water and the concentration was analyzed by NanoDrop on the ssDNA setting (Thermo Fisher Scientific). Probe sets with ssDNA concentration ranging from 200-500 ng/*µ*L, depending on the hairpin, were used for this study.

Fluorescent oligonucleotides were ordered from IDT with 5’ modification of either AlexaFluor 488, ATTO 550, ATTO 590, or ATTO 633. The sequences for fluorescent oligonucleotides and hairpins are included in Supplementary Files. Working dilutions of the hairpins (5 *µ*M) and tiling oligonucleotides (10 *µ*M) were made by diluting in IDTE pH 7.5 and stored at −20°C. Working dilutions of the fluorescent oligonucleotides (10 *µ*M) were made by diluting in UltraPure Water and stored at −20°C.

### Fluorescent oligonucleotide stripping for multiplexed Probe-Seq

Cells were incubated in 50% formamide solution (diluted in PBS) for 5 minutes at RT. The cells were then centrifuged at 2000 xg for 5 minutes at RT. The cells were resuspended in 1 mL of PBS and centrifuged again at 2000 xg for 5 minutes at RT. Hybridization of new fluorescent oligonucleotides was carried out as described above.

### Stain Index calculation

The Stain Index (SI) was calculated by measuring the geometric mean of the positive and negative populations as well as the standard deviation of the negative population using FlowJo Software. The SI was calculated as follows: (Geo. Mean_POS_ - Geo. Mean_NEG_) / (2 x SD_NEG_).

### Live cell RNA sequencing

*In vivo* retina electroporation was carried out as described previously at P2^10^. Two plasmids, CAG-BFP and Grik1^CRM4^-GFP, were electroporated simultaneously at a concentration of 1 *µ*g/*µ*L per plasmid. Retinas were harvested at P40. The electroporated and unelectroporated regions were processed separately. The electroporated region was dissociated as described above, and BFP^+^/GFP^+^ cells were FACS isolated into Trizol (Thermo Fisher Scientific, cat. #15596026). Cells from the unelectroporated region were used for Probe-Seq as described above. The RNA from the cells in Trizol was extracted following manufacturers protocol. The RNA-sequencing libraries were generated using the SMART-Seq v.4 and Nextera XT kits as described above.

### Histology and SABER FISH

For the mouse retina, adult CD1 mouse retinas were dissected and fixed in 4% PFA for 20 minutes at RT. The fixed retinas were cryoprotected in 30% sucrose (in PBS). Once submerged, the samples were embedded in 50%/15% OCT/Sucrose mixture in an ethanol/dry ice bath and stored at −80°C. The retinas were cryosectioned at 15 *µ*M thickness. For the human eye, formalin-fixed human postmortem eyes were obtained from Restore Life USA. Patient DRLU101818C was a 54-year-old male with no clinical eye diagnosis and the postmortem interval was 4 hours. Patient DRLU110118A was a 59-year-old female with no clinical eye diagnosis and the postmortem interval was 4 hours. A square (1 cm x 1 cm) of the human retina was cryoprotected, embedded, and cryosectioned as described above. For the *Drosophila* gut, the midgut was fixed in 4% PFA for 30 minutes at RT. For the chick retina, the central region of developing HH28 chick retina that contained the developing high acuity area was excised and fixed in 4% PFA for 20 minutes at RT. The retina was then cryoprotected, embedded as described above, and cryosectioned at 50 *µ*M thickness.

SABER FISH of retinal sections was carried out on Superfrost Plus slides (Thermo Fisher Scientific, cat. #1255015) using an adhesive hybridization chamber (Grace Bio-Labs, cat. #621502). For *Drosophila* guts, wholemount SABER FISH was performed in microcentrifuge tubes. For retinal sections, they were rehydrated with PBS for 5-10 minutes to remove the OCT on the slides. Subsequently, sections were completely dried to adhere the sections to the slides. Once dry, the adhesive chamber was placed to encase the sections. For both retinal sections and wholemount *Drosophila* guts, the samples were incubated in 0.1% PBS/Tween-20 (MilliporeSigma, cat. #P9416) for at least 10 minutes. The PBST was removed, and the samples were incubated with pre-warmed (43°C) 40% wHyb (2x SSC, 40% deionized formamide, 1% Tween-20, diluted in UltraPure Water) for at least 15 minutes at 43°C. The 40% wHyb was removed, and the samples were then incubated with 100 *µ*L of pre-warmed (43°C) Probe Mix (1 *µ*g of probe per gene, 96 *µ*L of Hyb1 solution (2.5x SSC, 50% deionized formamide, 12.5% Dextran Sulfate, 1.25% Tween-20), diluted up to 120 *µ*L with UltraPure Water) and incubated 16-48 hours at 43°C. The samples were washed twice with 40% wHyb (30 minutes/wash, 43°C), twice with 2x SSC (15 minutes/wash, 43°C), and twice with 0.1% PBST (5 minutes/wash, 37°C).

The samples were then incubated with 100 *µ*L of Fluorescent Oligonucleotide Mix (100 *µ*L of PBST, 2 *µ*L of each 10 *µ*M Fluorescent Oligonucleotide) for 15 minutes at 37°C. The samples were washed three times with PBST at 37°C for 5 minutes each and counterstained with DAPI (Thermo Fisher Scientific, cat. #D1306; 1:50,000 of 5 mg/mL stock solution in PBS) or WGA-405s (Biotium, cat. #290271; 1:100 of 1 mg/mL stock solution in PBS). Cell segmentation and cell calling algorithms were performed as described previously^24^.

### Imaging

Fluorescent images were acquired with W1 Yokogawa Spinning disk confocal microscope with 50 *µ*M pinhole disk and 488, 561, and 640 laser lines. The objectives used were either Plan Fluor 40x/1.3 or Plan Apo 60x/1.4 oil objectives, and the camera used was Andor Zyla 4.2 Plus sCMOS monochrome camera. Nikon Elements Acquisition Software (AR 5.02) was used for image acquisition and Fiji or Adobe Photoshop CS6 was used for image analysis. SABER FISH images were acquired as a z-stack and converted to a 2D image by maximum projection in Fiji.

### Data availability

Raw sequencing data and matrices of read counts for the mouse, chick, and *Drosophila* Probe-Seq are available at GEO: GSE135572.

### Code availability

All R scripts used for differential expression analysis are available in Supplementary Files.

## Supporting information

Oligonucleotide List

## Acknowledgments

We would like to thank former and current members of the Cepko and Tabin Labs for the insightful discussion and feedback. We thank P.M. Llopis, R. Stephansky, and the MicRoN core at Harvard Medical School for their assistance with microscopy. We thank C. Araneo, F. Lopez, and the Flow Cytometry Core Facility for their assistance with flow cytometry. We thank S. da Silva. for providing the image of *in situ* hybridization for FGF8 in the developing chick retina. We thank C. Cowan and B. Roska for providing the list of human retinal cell type-specific markers. This work was supported by the Howard Hughes Medical Institute (C.L.C. and N.P.), Edward R. and Anne G. Lefler Postdoctoral Fellowship (R.A.), and HSCI Internship Program (M.D.G).

## Author Contributions

R.A., E.R.W., S.W.L., and C.L.C. conceived the study and designed the experiments. R.A. executed the experiments and analyzed the data. M.G. contributed to the execution of the frozen nuclei protocol. E.R.W. performed the analysis of puncta quantification and stain index. J.C. provided the chick tissue. E.A.L. and N.P. provided the *Drosophila* tissue. R.A. and C.L.C. wrote the manuscript, and all authors edited the manuscript. C.L.C. supervised all aspects of the work.

## REFERENCES

1. Sanes, J.R. & Zipursky, S.L. Design principles of insect and vertebrate visual systems. Neuron 66, 15–36 (2010).

2. Vlasits, A.L., Euler, T. & Franke, K. Function first: classifying cell types and circuits of the retina. Current opinion in neurobiology 56, 8–15 (2019).

3. Macosko, E.Z. et al. Highly Parallel Genome-wide Expression Profiling of Individual Cells Using Nanoliter Droplets. Cell 161, 1202–1214 (2015).

4. Rheaume, B.A. et al. Single cell transcriptome profiling of retinal ganglion cells identifies cellular subtypes. Nature communications 9, 2759 (2018).

5. Shekhar, K. et al. Comprehensive Classification of Retinal Bipolar Neurons by Single-Cell Transcriptomics. Cell 166, 1308–1323 (2016).

6. Consortium, T. et al. Single-cell transcriptomics of 20 mouse organs creates a Tabula Muris. Nature 562, 367–372 (2018).

7. Han, X. et al. Mapping the Mouse Cell Atlas by Microwell-Seq. Cell 172, 1091–1107 (2018).

8. Regev, A. et al. The Human Cell Atlas. eLife 6 (2017).

9. Arlotta, P. et al. Neuronal subtype-specific genes that control corticospinal motor neuron development *in vivo*. Neuron 45, 207–221 (2005).

10. Matsuda, T. & Cepko, C.L. Controlled expression of transgenes introduced by *in vivo* electroporation. Proceedings of the National Academy of Sciences of the United States of America 104, 1027–1032 (2007).

11. Molyneaux, B.J. et al. DeCoN: genome-wide analysis of *in vivo* transcriptional dynamics during pyramidal neuron fate selection in neocortex. Neuron 85, 275–288 (2015).

12. Siegert, S. et al. Transcriptional code and disease map for adult retinal cell types. Nature neuroscience 15, 487–495 (2012).

13. Telley, L. et al. Sequential transcriptional waves direct the differentiation of newborn neurons in the mouse neocortex. Science (New York, N.Y.) 351, 1443–1446 (2016).

14. Xu, X. et al. Species and cell-type properties of classically defined human and rodent neurons and glia. eLife 7 (2018).

15. Jaitin, D.A. et al. Massively parallel single-cell RNA-seq for marker-free decomposition of tissues into cell types. Science (New York, N.Y.) 343, 776–779 (2014).

16. Klein, A.M. et al. Droplet barcoding for single-cell transcriptomics applied to embryonic stem cells. Cell 161, 1187–1201 (2015).

17. Picelli, S. et al. Smart-seq2 for sensitive full-length transcriptome profiling in single cells. Nature methods 10, 1096–1098 (2013).

18. Shalek, A.K. et al. Single-cell transcriptomics reveals bimodality in expression and splicing in immune cells. Nature 498, 236–240 (2013).

19. Amamoto, R. et al. FIN-Seq: Transcriptional profiling of specific cell types in frozen archived tissue from the human central nervous system. bioRxiv, 602847 (2019).

20. Hrvatin, S., Deng, F., O’Donnell, C.W., Gifford, D.K. & Melton, D.A. MARIS: method for analyzing RNA following intracellular sorting. PloS one 9 (2014).

21. Pan, Y., Ouyang, Z., Wong, W.H. & Baker, J.C. A new FACS approach isolates hESC derived endoderm using transcription factors. PloS one 6 (2011).

22. Pechhold, S. et al. Transcriptional analysis of intracytoplasmically stained, FACS-purified cells by high-throughput, quantitative nuclease protection. Nature biotechnology 27, 1038–1042 (2009).

23. Yamada, H. et al. Messenger RNA quantification after fluorescence activated cell sorting using intracellular antigens. Biochemical and Biophysical Research Communications 397, 425–428 (2010).

24. Kishi, J.Y. et al. SABER amplifies FISH: enhanced multiplexed imaging of RNA and DNA in cells and tissues. Nature methods 16, 533–544 (2019).

25. Klemm, S. et al. Transcriptional profiling of cells sorted by RNA abundance. Nature methods 11, 549–551 (2014).

26. Maeda, T. et al. Optimization of Recovery and Analysis of RNA in Sorted Cells in mRNA Quantification After Fluorescence-activated Cell Sorting. Annals of clinical and laboratory science 46, 571–577 (2016).

27. Yamada, H. et al. Messenger RNA quantification after fluorescence-activated cell sorting using *in situ* hybridization. Cytometry. Part A: the journal of the International Society for Analytical Cytology 77, 1032–1037 (2010).

28. Kishi, J.Y., Schaus, T.E., Gopalkrishnan, N., Xuan, F. & Yin, P. Programmable autonomous synthesis of single-stranded DNA. Nature chemistry 10, 155–164 (2018).

29. Sigurgeirsson, B., Emanuelsson, O. & Lundeberg, J. Sequencing degraded RNA addressed by 3’ tag counting. PloS one 9 (2014).

30. Krishnaswami, S.R. et al. Using single nuclei for RNA-seq to capture the transcriptome of postmortem neurons. Nature protocols 11, 499–524 (2016).

31. Lake, B.B. et al. Neuronal subtypes and diversity revealed by single-nucleus RNA sequencing of the human brain. Science (New York, N.Y.) 352, 1586–1590 (2016).

32. Cowan, C.S. et al. Cell types of the human retina and its organoids at single-cell resolution: developmental convergence, transcriptomic identity, and disease map. bioRxiv, 703348 (2019).

33. Dutta, D. et al. Regional Cell-Specific Transcriptome Mapping Reveals Regulatory Complexity in the Adult Drosophila Midgut. Cell reports 12, 346–358 (2015).

34. Hung, R.-J. et al. A cell atlas of the adult Drosophila midgut. bioRxiv, 410423 (2018).

35. Doup, D.P., Marshall, O.J., Dayton, H., Brand, A.H. & Perrimon, N. Drosophila intestinal stem and progenitor cells are major sources and regulators of homeostatic niche signals. Proceedings of the National Academy of Sciences of the United States of America 115, 12218–12223 (2018).

36. Bruhn, S.L. & Cepko, C.L. Development of the pattern of photoreceptors in the chick retina. The Journal of neuroscience: the official journal of the Society for Neuroscience 16, 1430–1439 (1996).

37. da Silva, S. & Cepko, C.L. Fgf8 Expression and Degradation of Retinoic Acid Are Required for Patterning a High-Acuity Area in the Retina. Developmental cell 42, 68–81 (2017).

38. Eastburn, D.J., Sciambi, A. & Abate, A.R. Identification and genetic analysis of cancer cells with PCR-activated cell sorting. Nucleic acids research 42 (2014).

39. Pellegrino, M. et al. RNA-Seq following PCR-based sorting reveals rare cell transcriptional signatures. BMC genomics 17, 361 (2016).

40. Rouhanifard, S.H. et al. ClampFISH detects individual nucleic acid molecules using click chemistry-based amplification. Nature biotechnology 37, 84–89 (2018).

41. Wang, F. et al. RNAscope: a novel *in situ* RNA analysis platform for formalin-fixed, paraffin-embedded tissues. The Journal of molecular diagnostics: JMD 14, 22–29 (2012).

42. Alles, J. et al. Cell fixation and preservation for droplet-based single-cell transcriptomics. BMC biology 15, 44 (2017).

43. Attar, M. et al. A practical solution for preserving single cells for RNA sequencing. Scientific reports 8, 2151 (2018).

44. Chen, J. et al. PBMC fixation and processing for Chromium single-cell RNA sequencing. Journal of translational medicine 16, 198 (2018).

45. Hamburger, V. & Hamilton, H.L. A series of normal stages in the development of the chick embryo. Journal of morphology 88, 49–92 (1951).

46. Aken, B.L. et al. Ensembl 2017. Nucleic acids research 45 (2017).

47. Dobin, A. et al. STAR: ultrafast universal RNA-seq aligner. Bioinformatics (Oxford, England) 29, 15–21 (2013).

48. Okonechnikov, K., Conesa, A. & Garca-Alcalde, F. Qualimap 2: advanced multi-sample quality control for high-throughput sequencing data. Bioinformatics (Oxford, England) 32, 292–294 (2016).

49. Anders, S., Pyl, P.T. & Huber, W. HTSeq–a Python framework to work with high-throughput sequencing data. Bioinformatics (Oxford, England) 31, 166–169 (2015).

50. Love, M.I., Huber, W. & Anders, S. Moderated estimation of fold change and dispersion for RNA-seq data with DESeq2. Genome biology 15, 550 (2014).

51. Subramanian, A. et al. Gene set enrichment analysis: a knowledge-based approach for interpreting genome-wide expression profiles. Proceedings of the National Academy of Sciences of the United States of America 102, 15545–15550 (2005).

