## Supplementary Figures for "Probe-Seq enables transcriptional profiling of specific cell types from heterogeneous tissue by RNA-based isolation"

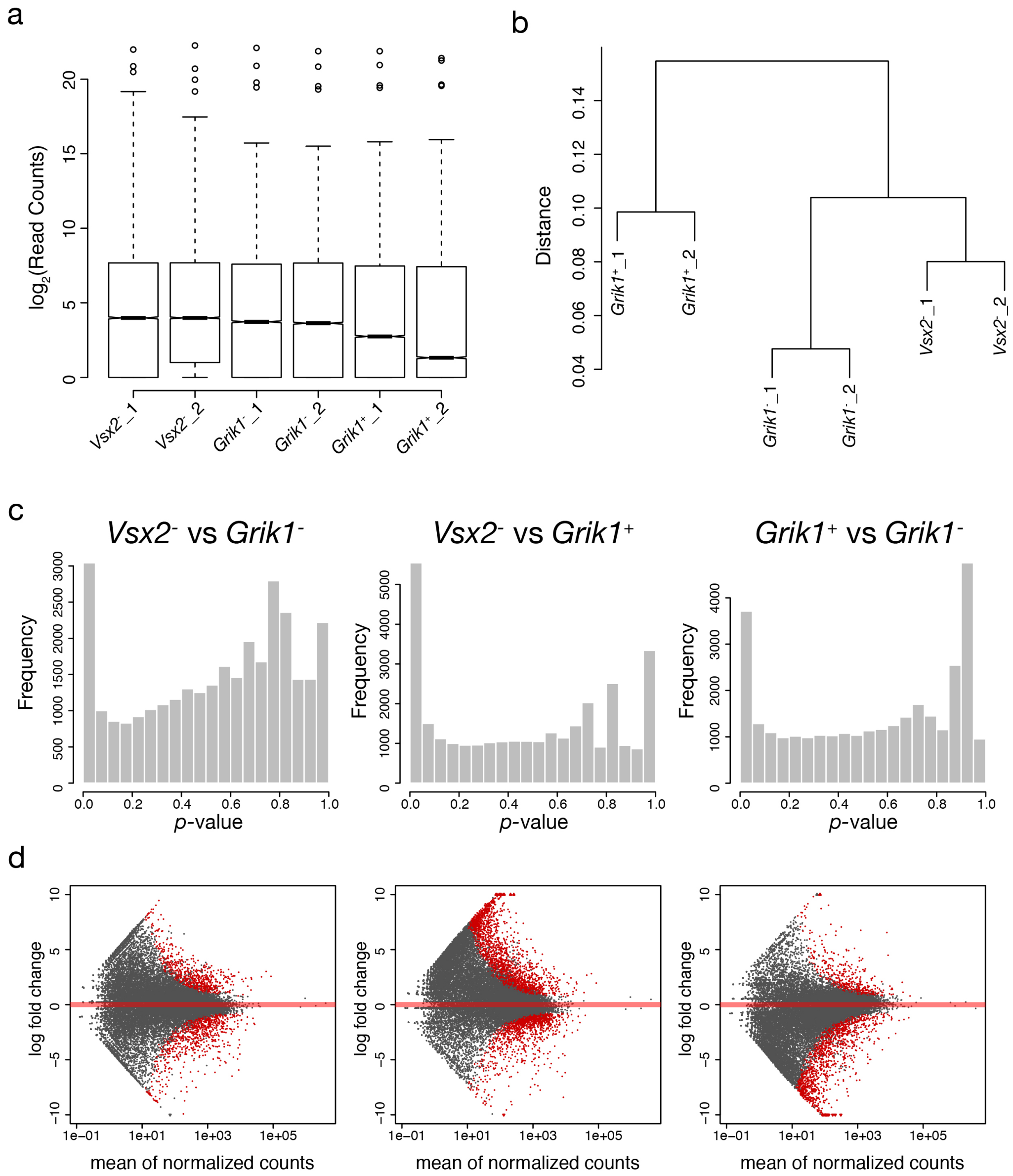


**Supplementary Figure 1: Quality control of mouse Probe-Seq RNA sequencing.**

a) $Log_2$-transformed read distribution plot for sequenced mouse *Vsx2/Grik1* Probe-Seq samples. b) Dendrogram of read counts shows clustering of *Vsx2*$^-$, *Grik1*$^-$, and *Grik1*$^+$ samples. c) Plots of frequencies of *p*-values shows an even distribution of null *p*-values. d) MA plots of $log_2$ fold change vs. mean of normalized counts. Red dots indicate genes that are differentially expressed.

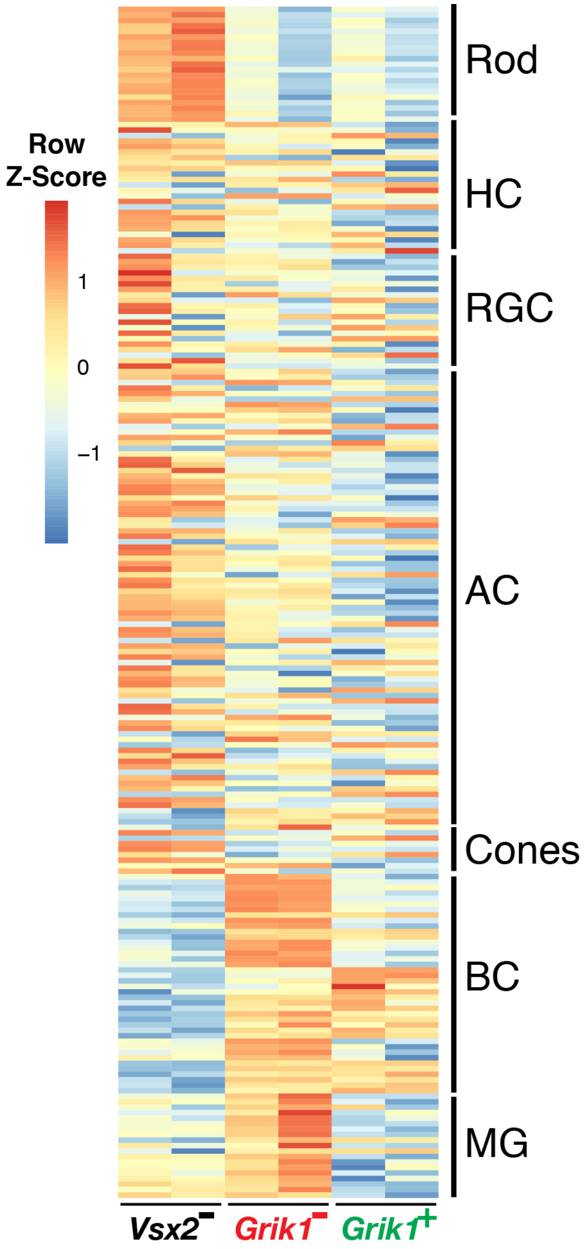


**Supplementary Figure 2: BC and MG markers are enriched in the *Grik*$^{-}$ and *Grik1*$^{+}$ populations.**

A heatmap representing relative expression levels of major retinal cell class markers previously identified by scRNA sequencing[3] that are differentially expressed (adjusted *p*-value<0.05) between *Vsx2*$^{-}$, *Grik1*$^{-}$, and *Grik1*$^{+}$ populations.

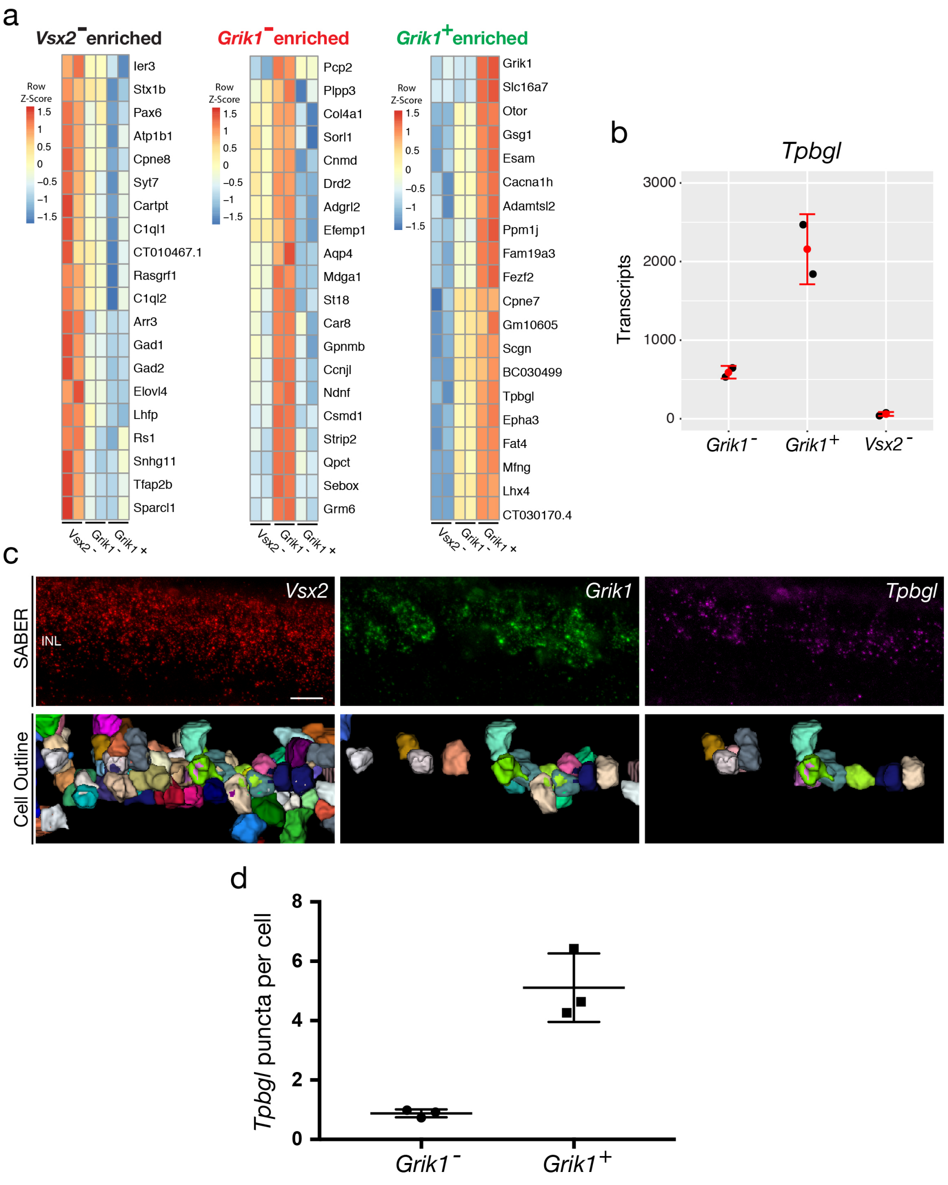


**Supplementary Figure 3: Validation of a *Grik1*$^{+}$ population-enriched transcript, *Tpbgl,* by SABER FISH.**

a) Heatmaps of unbiased top 20 enriched genes for each population (*Vsx2*$^{-}$, left; *Grik1*$^{-}$, middle; *Grik1*$^{+}$, right). b) Quantification of *Tpbgl* transcripts in *Vsx2*$^{-}$, *Grik1*$^{-}$, *Grik1*$^{+}$ populations based on Probe-Seq. c) Images of a mouse retinal section following the SABER FISH protocol using gene-specific probe sets for *Vsx2, Grik1,* and *Tpbgl*. Reconstruction of cell outlines was carried out using the cell segmentation algorithm described previously[24]. d) Quantification of *Tpbgl* puncta in *Grik1*$^{-}$ cells vs *Grik1*$^{+}$ cells.

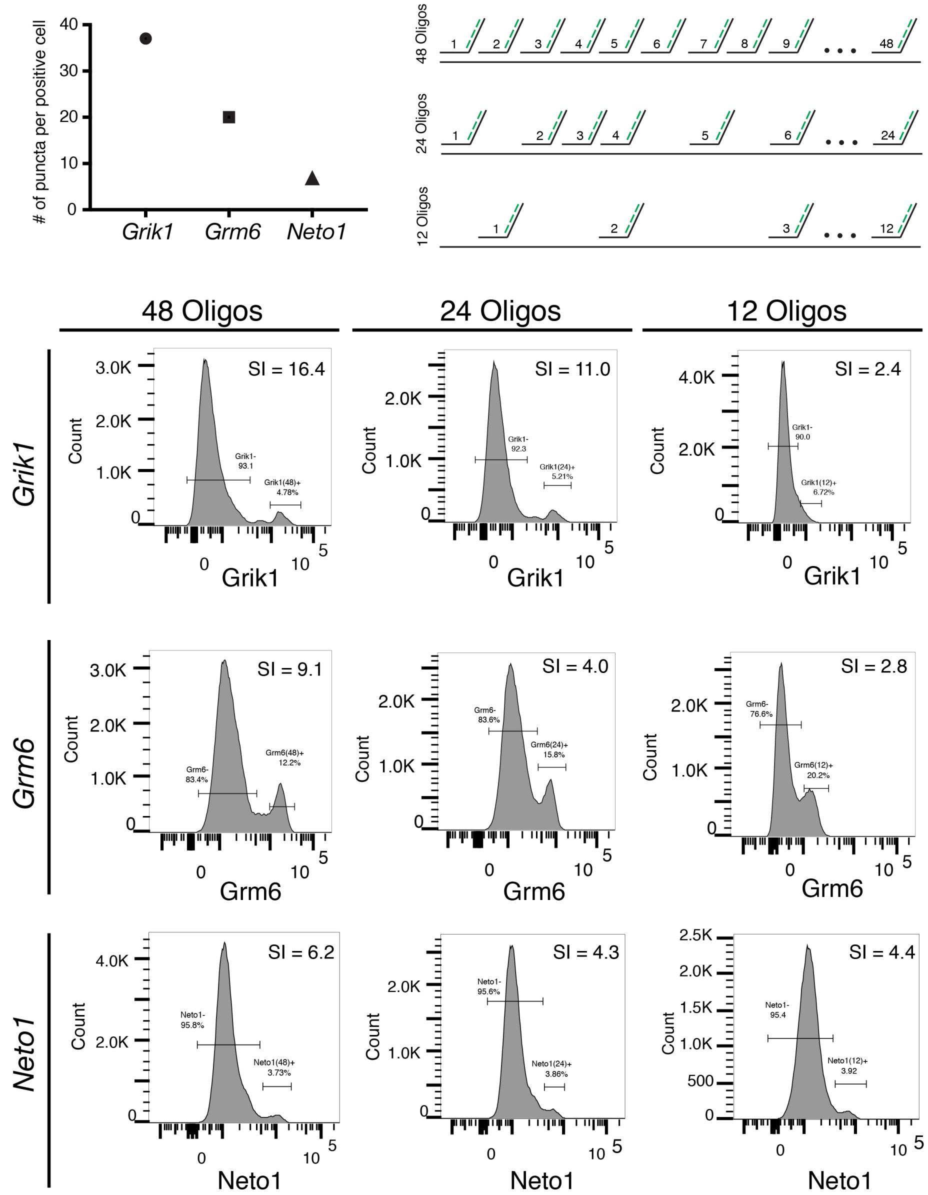


**Supplementary Figure 4: Probe-Seq stain index is correlated with the number of tiling oligonucleotides and the expression level.**

a) Quantification of number of puncta per positive cell for *Grik1, Grm6,* and *Neto1* in the mouse retina. b) Schematic of the experiment. For each gene, 48, 24, or 12 randomly-chosen tiling oligonucleotides were pooled for gene-specific probe set synthesis. c) Flow cytometry histograms of *Grik1* (upper row)*, Grm6* (middle row)*,* and *Neto1* (bottom row) with either 48 (left column), 24 (middle column), and 12 (right column) tiling oligonucleotides. SI, Stain Index.

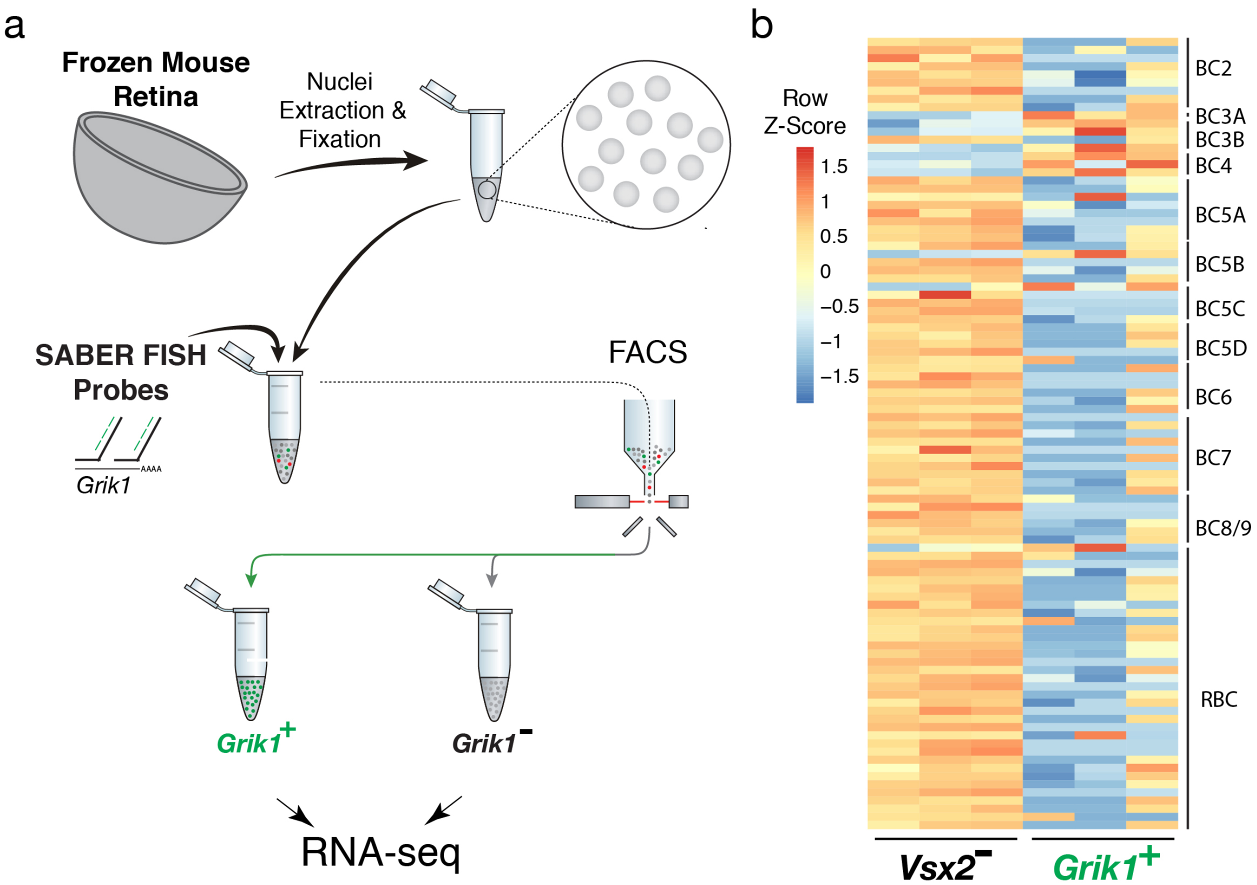


**Supplementary Figure 5: Isolation and transcriptional profiling of specific BC subtypes from the frozen mouse retina.**

a) Schematic of Probe-Seq for the fresh frozen adult mouse retina. Single nuclei were isolated and fixed. Nuclei were incubated with a SABER FISH probe set for *Grik1* and then incubated with fluorescent oligonucleotides. *Grik1*$^+$ and *Grik1*$^-$ populations were isolated by FACS for downstream RNA sequencing. b) A heatmap representing relative expression levels of mouse BC subtype markers previously identified by scRNA sequencing[5] that are differentially expressed (adjusted *p*-value<0.05) between *Grik1*$^-$, and *Grik1*$^+$ populations.

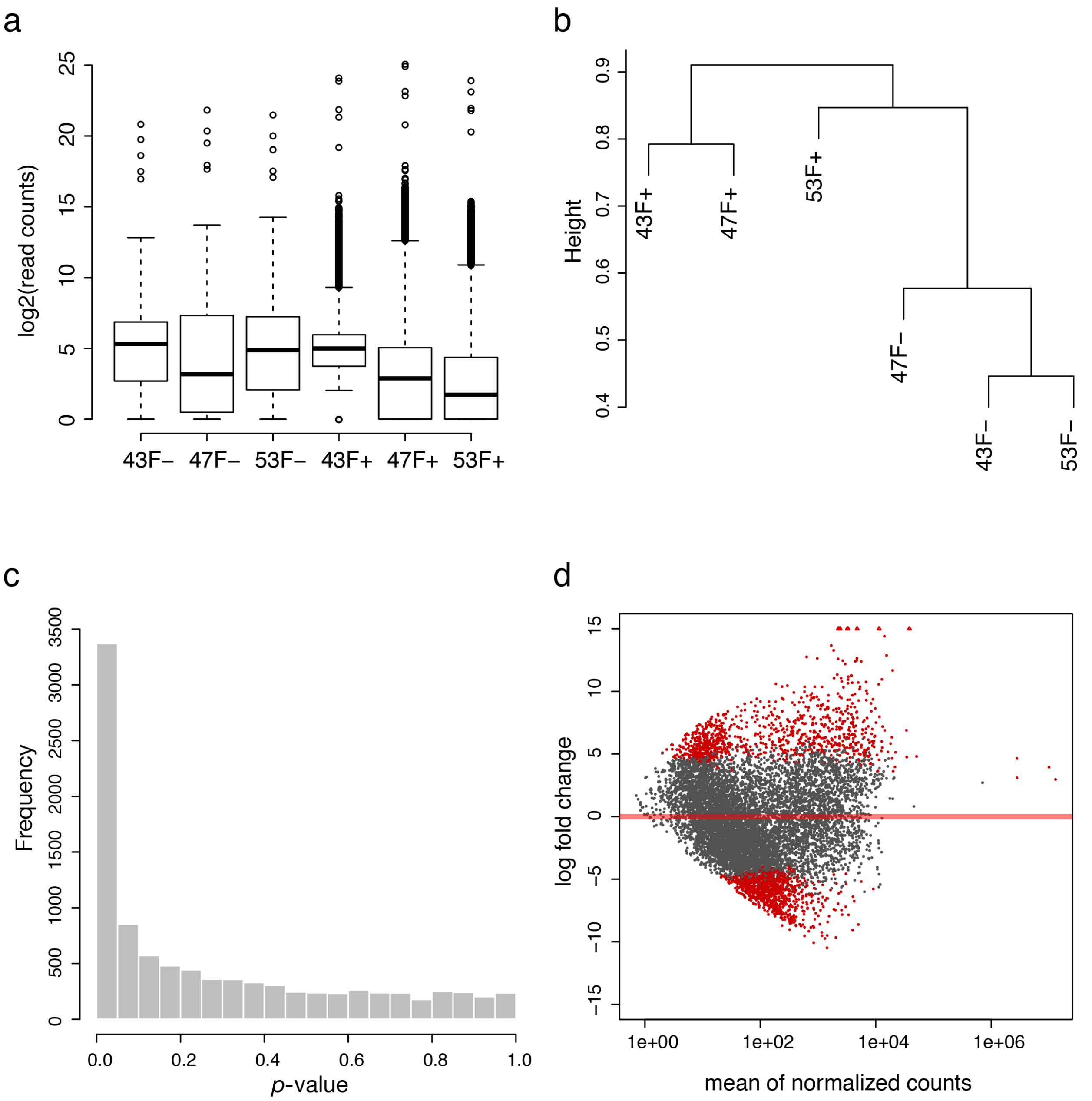


**Supplementary Figure 6: Quality control of human nuclear Probe-Seq RNA sequencing.**

a) $Log_2$-transformed read distribution plot for sequenced human *GRM6* Probe-Seq samples. b) Dendrogram of read counts shows clustering of *GRM6*$^-$ *and GRM6*$^+$ samples. c) Plot of frequencies of *p*-values shows an even distribution of null *p*-values. d) MA plot of $log_2$ fold change vs. mean of normalized counts. Red dots indicate genes that are differentially expressed between *GRM6*$^+$ and *GRM6*$^-$ samples.

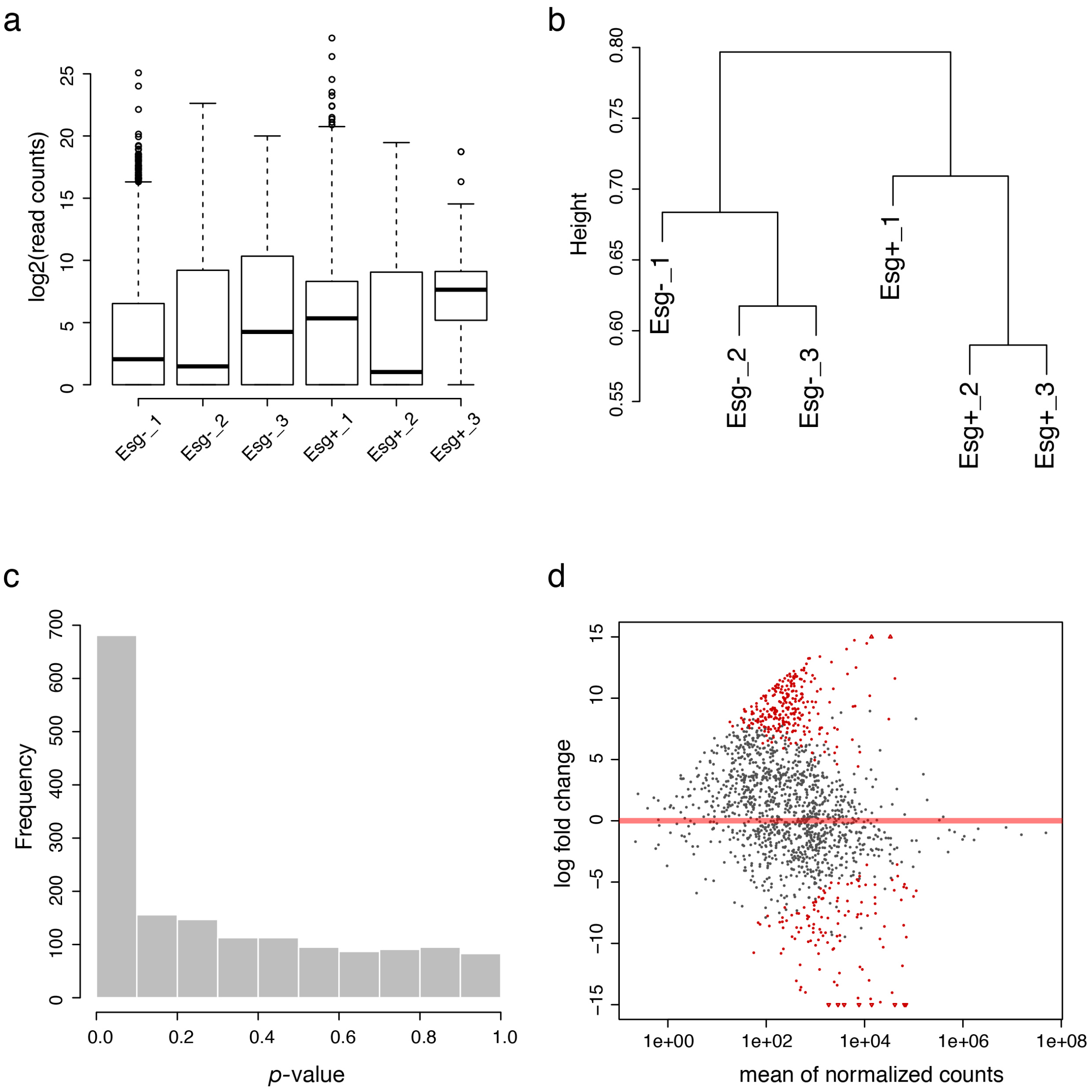


**Supplementary Figure 7: Quality control of *Drosophila* Probe-Seq RNA sequencing.**

a) $Log_2$-transformed read distribution plot for sequenced *Drosophila esg* Probe-Seq samples. b) Dendrogram of read counts shows clustering of *esg*$^-$ and *esg*$^+$ samples. c) Plot of frequencies of *p*-values shows an even distribution of null *p*-values. d) MA plot of $log_2$ fold change vs. mean of normalized counts. Red dots indicate genes that are differentially expressed.

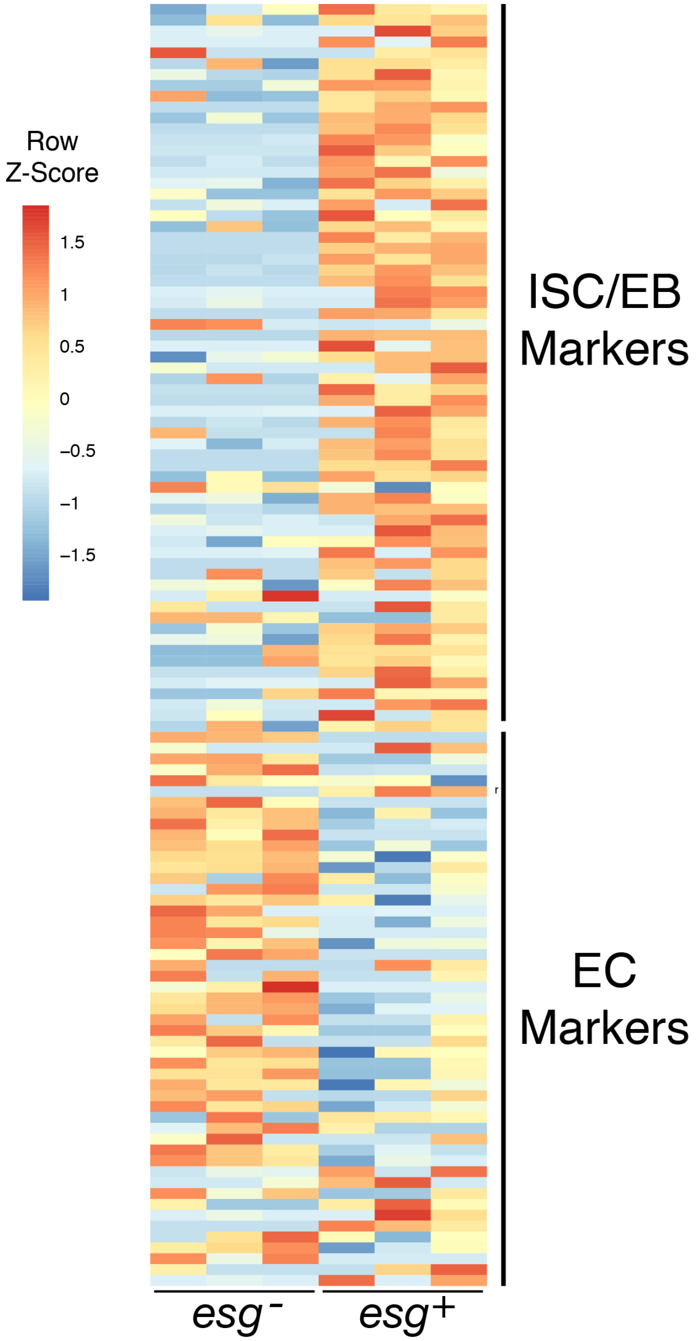


**Supplementary Figure 8: Heatmap of ISC/EB and EC markers based on DamID transcriptional profiling.**

A heatmap representing relative expression levels of *Drosophila* ISC/EB and EC markers previously identified by DamID transcriptional profiling (adjusted *p*-value<0.05) between *esg*$^{-}$, and *esg*$^{+}$ populations[35]. ISC, Intestinal Stem Cell; EB, Enteroblast; EC, Enterocyte.

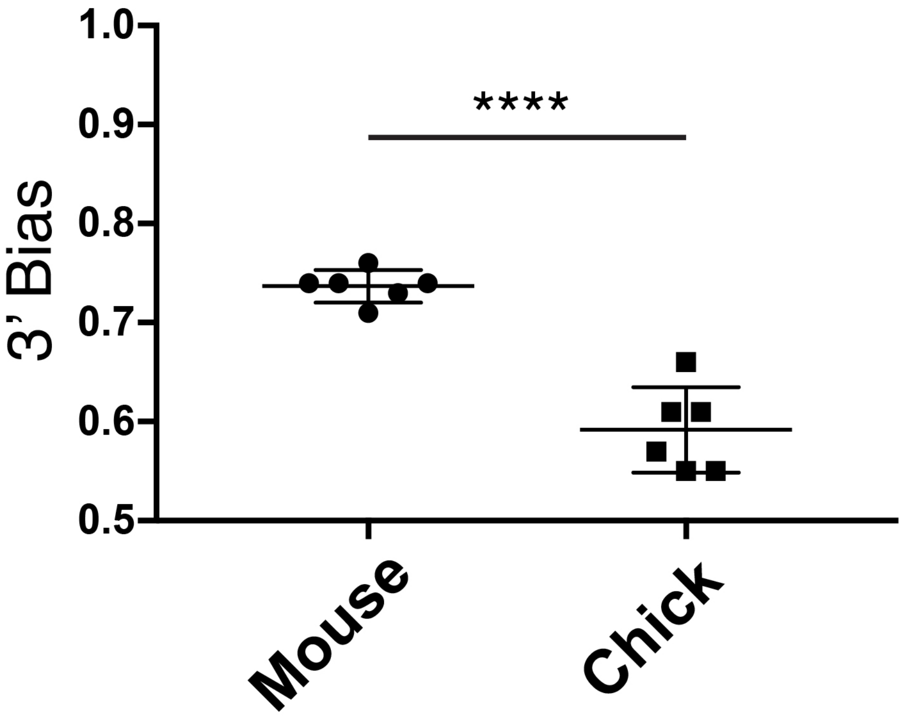


**Supplementary Figure 9: 3' bias is significantly decreased for chick Probe-Seq RNA preparation compared to that of mouse.**

Quantification of 3' bias from adult mouse *Vsx2/Grik1* Probe-Seq reads and developing chick *FGF8* Probe-Seq reads. All results are expressed as the mean ± SD. **** $p < 0.0001$. Unpaired Student's t-test.

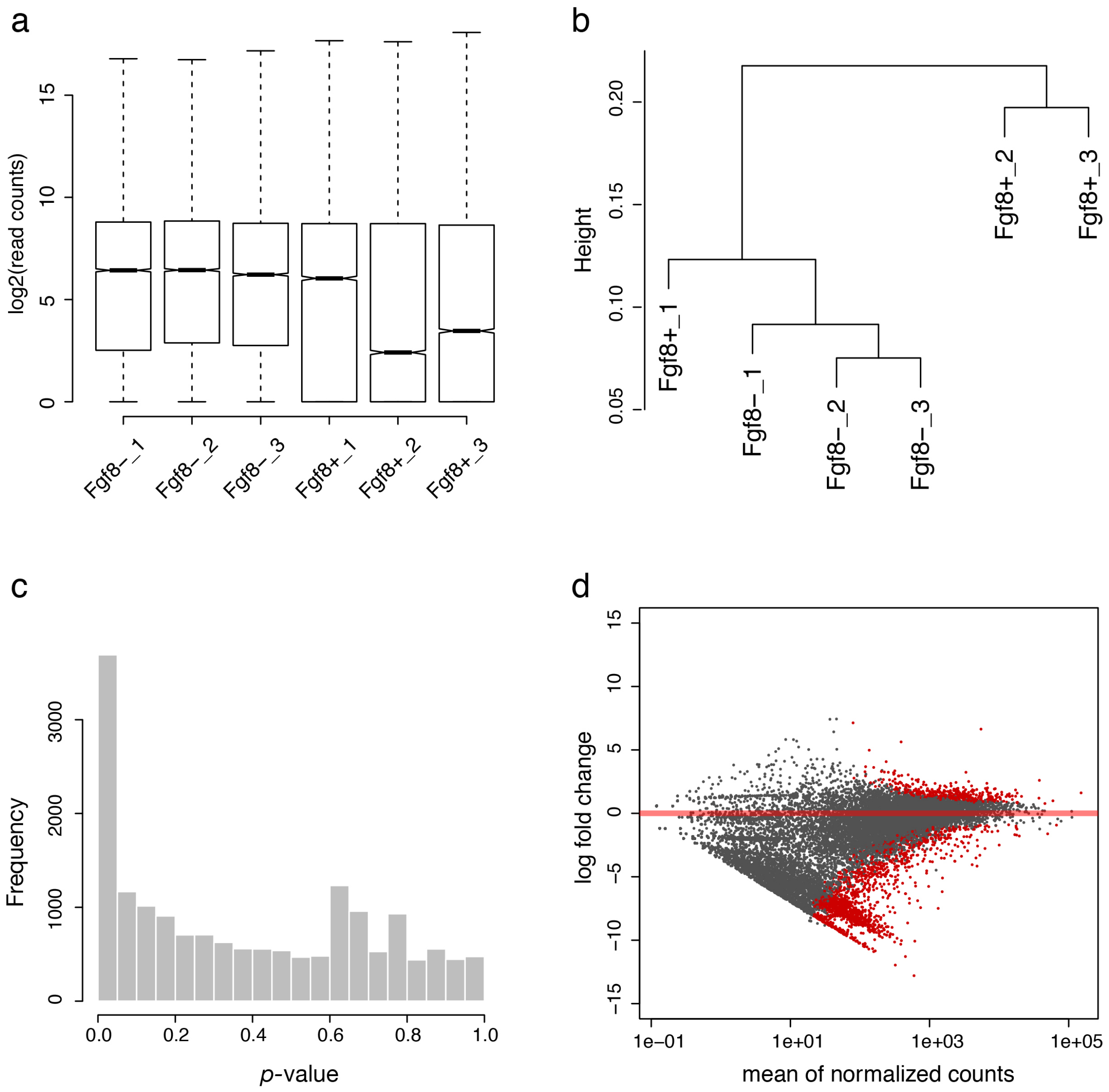


**Supplementary Figure 10: Quality control of developing chick retina Probe-Seq RNA sequencing.**

a) Log$_2$-transformed read distribution plot for sequenced chick Probe-Seq samples. b) Dendrogram of read counts shows clustering of *FGF8*$^{-}$ and *FGF8*$^{+}$ samples except FGF8$^{+}$-1. c) Plot of frequencies of *p*-values shows an even distribution of null *p*-values. d) MA plot of log$_2$ fold change vs. mean of normalized counts. Red dots indicate genes that are differentially expressed.

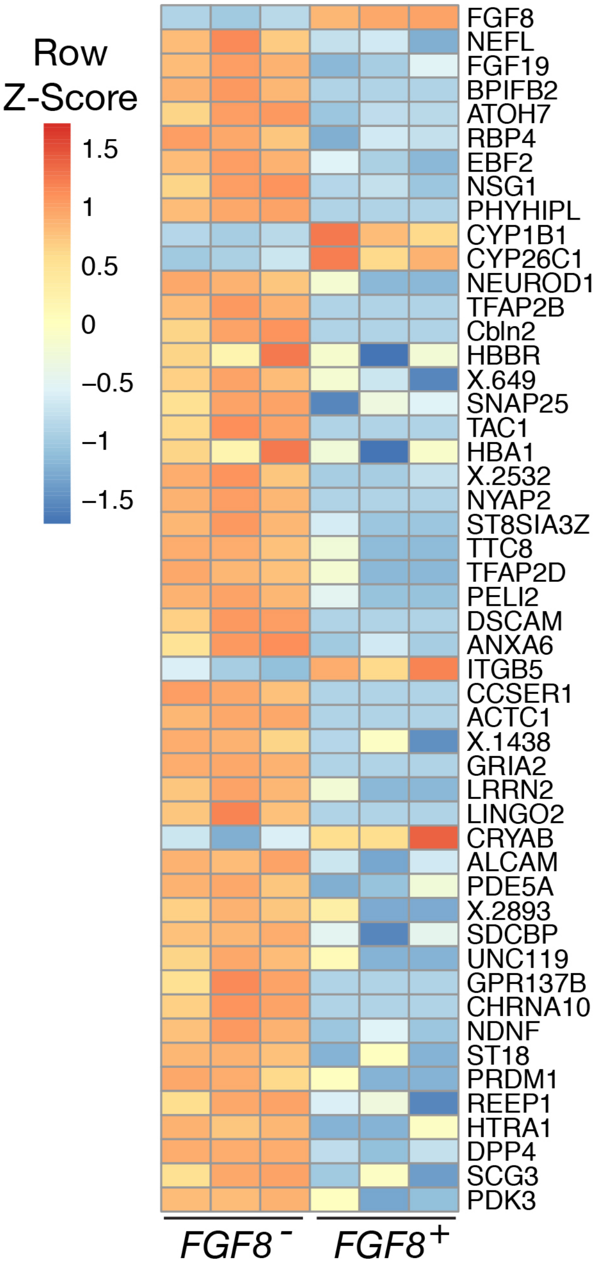


**Supplementary Figure 11: Early differentiation markers are enriched in the *FGF8*$^{-}$ population.**

A heatmap of top 50 unbiased differentially expressed genes between *FGF8*$^{+}$ and *FGF8*$^{-}$ populations in the developing chick retina.
